# Weak brain-mental health associations in population neuroimaging tasks reflect a signal evocation bottleneck

**DOI:** 10.1101/2025.06.11.659091

**Authors:** Cole Korponay, Julia E. Cohen-Gilbert, Poornima Kumar, Nathaniel G. Harnett, Adrian A. Medina, You Cheng, Brent P. Forester, Kerry J. Ressler, Jure Demsar, Blaise B. Frederick, Christian F. Beckmann, David G. Harper, Lisa D. Nickerson

## Abstract

Large-scale neuroimaging studies best powered to investigate mental health biomarkers continue to administer a set of in-scanner tasks that, to date, yield weak brain–mental health associations. Triangulating the source of these weak associations is critical to directing the focus of future work. Using these tasks’ strong brain–cognition associations as a positive control, we analyzed potential sources of weak brain-mental health associations across five popular fMRI tasks and dozens of task-evoked brain networks. Across tasks and networks, latent mental health factors showed substantially weaker brain associations than cognition despite comparable measurement reliability and greater interindividual variability. Moreover, increasing brain-feature dimensionality improved cognitive but not mental health signal, and task-evoked network profiles were 1.8–2.3 times more homogeneous and 3.5 times lower-dimensional than mental health profiles. Findings implicate a signal evocation bottleneck: common fMRI tasks elicit little interindividual variation in mental health–related brain activity relative to the diversity of mental health itself. Overall, these findings motivate a shift in focus to the development and use of tasks in large-scale studies more explicitly designed to evoke mental health-relevant brain activity.

**Significance Statement:** Identification of robust individual-specific mental health-related brain signals from fMRI remains elusive, limiting the clinical utility of brain imaging. To investigate why, we systematically evaluated brain signals evoked by tasks used by the Human Connectome Project-Young Adult study, which are now commonly included in many large-scale neuroimaging studies. Our findings suggest that, more so than analytical or measurement constraints on detecting these signals, a primary limitation is a failure of these prevalent fMRI tasks to evoke mental health-relevant brain signals in the first place. These findings suggest that progress in psychiatric neuroimaging may require tasks more explicitly designed to evoke mental health-relevant brain activity.

## Introduction

Our experiences of and responses to cognitive challenges, positive and negative emotions, and social interactions are core determinants and reflections of our mental health. Functional magnetic resonance imaging (fMRI) has become a central tool for probing the role of neurobiology in these clinically relevant domains^1–4^, formalized in the Research Domain Criteria (RDoC) framework^5,6^. RDoC is predicated on considering mental illness in terms of deviations from normal neurobehavioral functions across these domains (e.g., cognitive system (CS), positive valence (PVS) and negative valence (NVS) systems). The proliferation of large-scale fMRI datasets over the past decade, which have included tasks specifically selected to probe these functional domains, has paralleled optimism that fMRI-based brain features (e.g., functional connectivity, task activation) could yield meaningful biomarkers of psychiatric illness. Yet, a persistent empirical pattern has emerged: across datasets, fMRI task conditions, and analytic approaches, fMRI-based brain features display moderately strong associations with cognitive function but relatively weak associations with non-cognitive domains of mental health, such as internalizing or well-being^1,4,7–24^. Even brain activity during tasks like the emotional face matching task – commonly used to index interindividual variability in function related to the NVS – has recently been found to predominantly reflect function related to the CS^14,25,26^. This has contributed to limited clinical utility of fMRI in psychiatry to date.

While small sample sizes were thought to be the main limitation to identifying brain-based mental health markers, analyses of new large-scale datasets show that this is not the case^1,4,7–24^. This has led to deeper investigations into why fMRI brain-based markers of mental health have not been realized. One potential limitation relates to the quality of behavioural data^1,7^. Compared to measures of cognition – which tend to be performance-based and relatively stable over time – non-cognitive mental health measures are primarily based on self-report or subjective evaluations that are more variable over time. As such, these non-cognitive measures typically display relatively low reliability^27^. Recent work demonstrates that the reliability of behavioral phenotypes is an important moderator of brain-behavior relationship strengths^28^.

Analytic techniques such as hierarchical factor analysis (hFA)^29^, which derive latent, composite scores with higher reliability by aggregating across many individual assessments^30,31^, offer one possible solution for this issue. Non-cognitive mental health is also often assumed to be more range-restricted than cognition in normative samples characteristic of many large-scale neuroimaging studies, limiting the ability to capture interindividual variance. Other potential limiting factors relate to fMRI data quality and analytical approaches. Optimization efforts in these domains have included the development of more rigorous fMRI signal denoising methods^32^, more sophisticated brain feature modeling^13,33,34^ paired with model validation techniques^35,36^, and the curation of larger and/or more deeply sampled datasets^37^. Collectively, these approaches share the underlying assumption that mental health-relevant brain signal is present in available task fMRI data, but remains difficult to detect under current measurement and modeling constraints. Advances in addressing these mental health signal detection constraints have yielded unmistakeable gains in brain-behaviour/mental health effect sizes and replicability, though effect sizes remain marginal.

In addition to limitations in signal *detection*, the dearth of clinically useful fMRI-based biomarkers of psychopathology could also be due to a limited ability of prevailing fMRI tasks to *evoke* strong mental health-relevant brain signals. While there exist a multitude of fMRI tasks, most large-scale neuroimaging studies with sufficient power to examine brain-behaviour/mental health relationships draw from a circumscribed set of tasks originally designed to localize functional processes in the brain^38^. The adoption of this overlapping set of tasks across large-scale studies was meant to facilitate harmonization and combination across datasets. However, given the original intent of such tasks to minimize individual differences for the sake of maximizing functional localization signal, these tasks involve relatively simple task demands and blocked designs that make them scalable and suitable for a range of populations but possibly limited for assessing interindividual variability. Possibly in relation to their simplicity, recent work even demonstrates that subject arousal levels dwindle during such classic fMRI tasks^39^.

Though these tasks – as well as more recent movie-watching tasks – evoke brain activity with stronger links to behavior than resting-state conditions, they appear to have similarly limited capacity to elicit mental health-relevant interindividual variation in brain activity^7,18,40^.

Disentangling the impacts of signal detection and signal evocation bottlenecks is critical for informing the direction of future work. In particular, advances in mitigating signal detection constraints may offer limited returns if there is a fundamental signal evocation issue with common task fMRI paradigms. To investigate, we leveraged data from five common fMRI tasks used in many large-scale neuroimaging cohorts, spanning affective, motivational, social-cognitive, and cognitive-control domains (emotion, gambling, social cognition, working memory, and movie-watching) and 87 behavioral and neuropsychiatric measures of cognitive and non-cognitive mental health in the Human Connectome Project–Young Adult (HCP-YA) cohort (n=1200). In addition to rich task fMRI and behavioral data available in HCP-YA, it comprises a normative range of mental health spanning generally healthy individuals to those with subclinical depression and anxiety and moderate-to-heavy alcohol and cannabis users. Since RDoC casts mental illness as deviations from normative biobehavioral functioning, it is important to map variability in large-scale normative populations like HCP-YA. If the task batteries in these normative large-scale studies are ineffective at eliciting mental health-relevant brain activity to assess normative mental health heterogeneity, this creates a challenge for identifying the deviations that define mental illness using these datasets.

In modeling the HCP-YA task fMRI data, given that the cognition versus non-cognition brain signal discrepancy is observed across fMRI-based brain features (e.g., regional activation, functional connectivity, dynamics, functional topography), we prioritized feature maximization of subject discriminability, across-sample reproducibility, and neurobiological interpretability at the scale of distributed systems. Prior work shows that large-scale network–level properties yield higher subject discriminability^20^ and stronger reproducibility^41^ than fine-grained edge-level features, and RDoC-relevant functions (e.g. cognitive, emotional, and social processes) are increasingly understood to be subserved by distributed brain networks rather than localized regions or single connections^42–45^. In the context of task and movie-watching fMRI, in which stimuli are time-locked across individuals, tensor independent component analysis (tICA) provides a robust, data-driven method to identify and quantify the engagement of such large-scale networks across individuals^25,46^. Unlike traditional group-ICA followed by dual regression — which projects individuals onto static network spatial maps — tICA decomposes the subject × time × space tensor simultaneously, preserving individual differences in both the spatial and temporal dimensions and yielding subject loadings on each component that may more sensitively capture interindividual variation in network engagement. Moreover, because tICA is applied to task timeseries, the resulting components specifically reflect task-evoked networks, rather than canonical time-averaged resting-state networks that may not capture the primary loci of task-evoked individual differences. Together, these properties make tICA particularly well-suited for brain-behavior analyses targeting distributed system-level individual differences in cognitive and non-cognitive aspects of mental health.

Thus, we applied tICA to the HCP-YA task fMRI data to identify and compute individual subject loadings on evoked network states, and applied Schottner and colleague’s hierarchal factor analysis (hFA)^29^ model to 87 behavioral and neuropsychiatric assessments to compute high-reliability mental health factor scores in five latent domains: cognition (reflecting the CS), positive affect/well-being (reflecting the PVS), negative affect/internalizing (reflecting the NVS), processing speed, and substance use. Subsequently, we quantified univariate and multivariate relationships between the brain and mental health data to benchmark the magnitude of cognitive versus non-cognitive mental health signal disparity for each fMRI task. Next, we systematically characterized the latent structure and measurement properties of the brain and mental health data to examine potential sources of low non-cognitive mental health signal in the task fMRI data. Finally, to examine the consequences of the low mental health signal we derived biotypes independently from the task-fMRI and mental health data and tested their correspondence and relative informativeness.

## Results

### Task-Evoked Brain Networks

tICA identified 77 evoked large-scale network states across five fMRI paradigms (Emotion, Gambling, Working Memory, Social Cognition, Movie-Watching) in the split-half discovery sample (Group 1) (**Fig. 1**).

**Figure 1.**
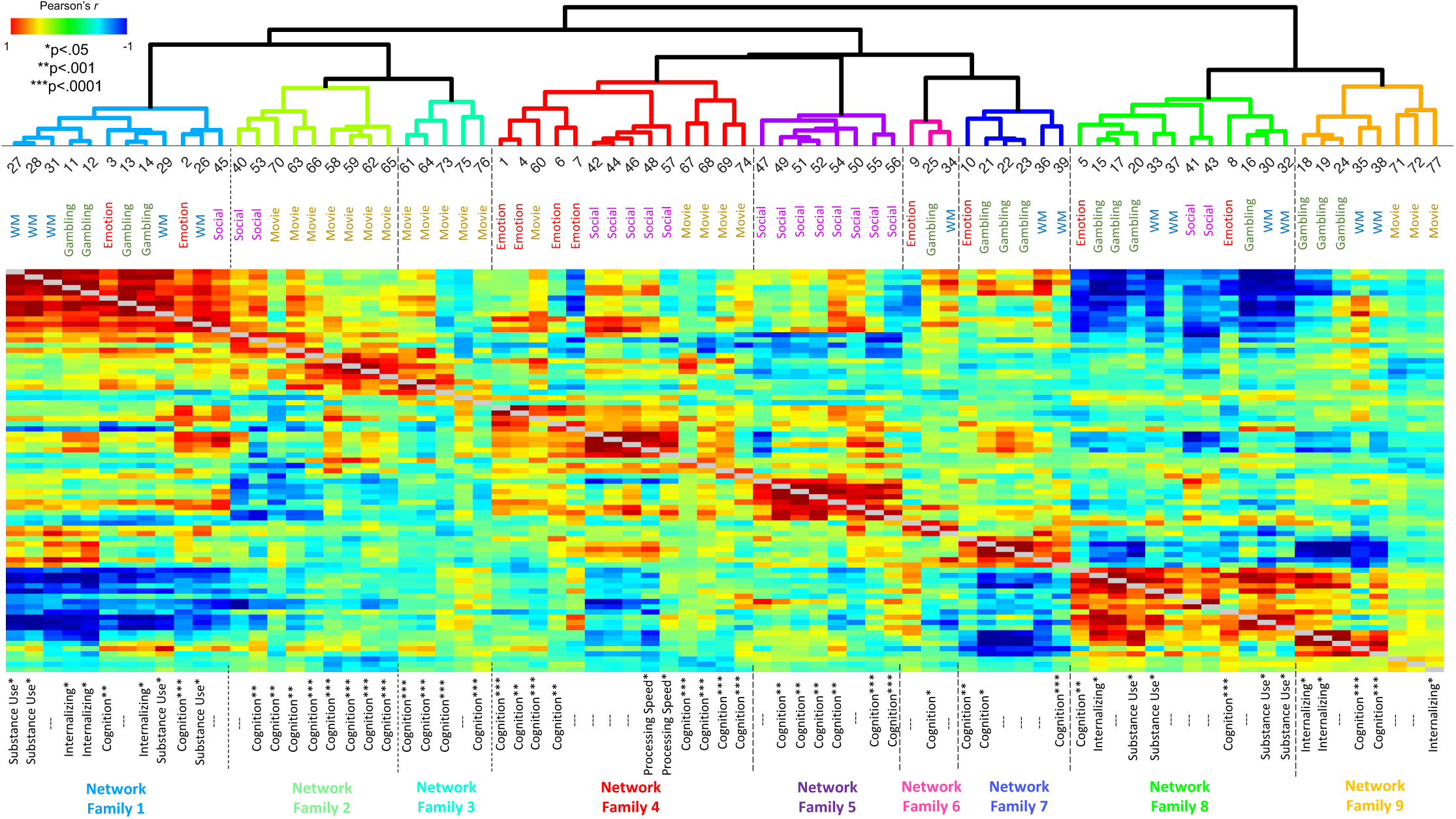
Hierarchical clustering of task-evoked network spatial maps. The heat map depicts the correlation matrix of unthresholded network spatial maps, with red indicating positively correlated spatial maps and blue indicating negatively correlated spatial maps. Task labels above each column in the heat map show the task each network was evoked by (i.e., Emotion, Gambling, Working Memory, Social Cognition, or Movie-Watching) and Factor labels below the heat map show whether cross-subject variability in the network’s activation during the specific task correlated with variability in one of the five mental health domains: Cognition, Internalizing/negative affect, Well-being/ positive affect, Processing Speed, and Substance Use (*p<.05 uncorrected; **p<.001 uncorrected; ***p<.0001 Bonferroni corrected threshold for 385 tests (77 networks * 5 mental health factors)).

Individual network state maps have been made viewable and accessible in a “Task-Evoked Network Atlas” (https://identifiers.org/neurovault.collection:20820). tICA-resolved network maps reflect the joint engagement of distinct sets of canonical networks during different task demands **(Fig. 2)**. Hierarchical clustering on the 77-network state similarity matrix identified nine Network Families (**Figs. 1-2**), consistent with the notion of a circumscribed set of canonical large-scale networks that manifest in context-dependent spatial variants^47^ and undergo reconfigurations in network-network coupling^48^ to meet particular task demands.

**Figure 2.**
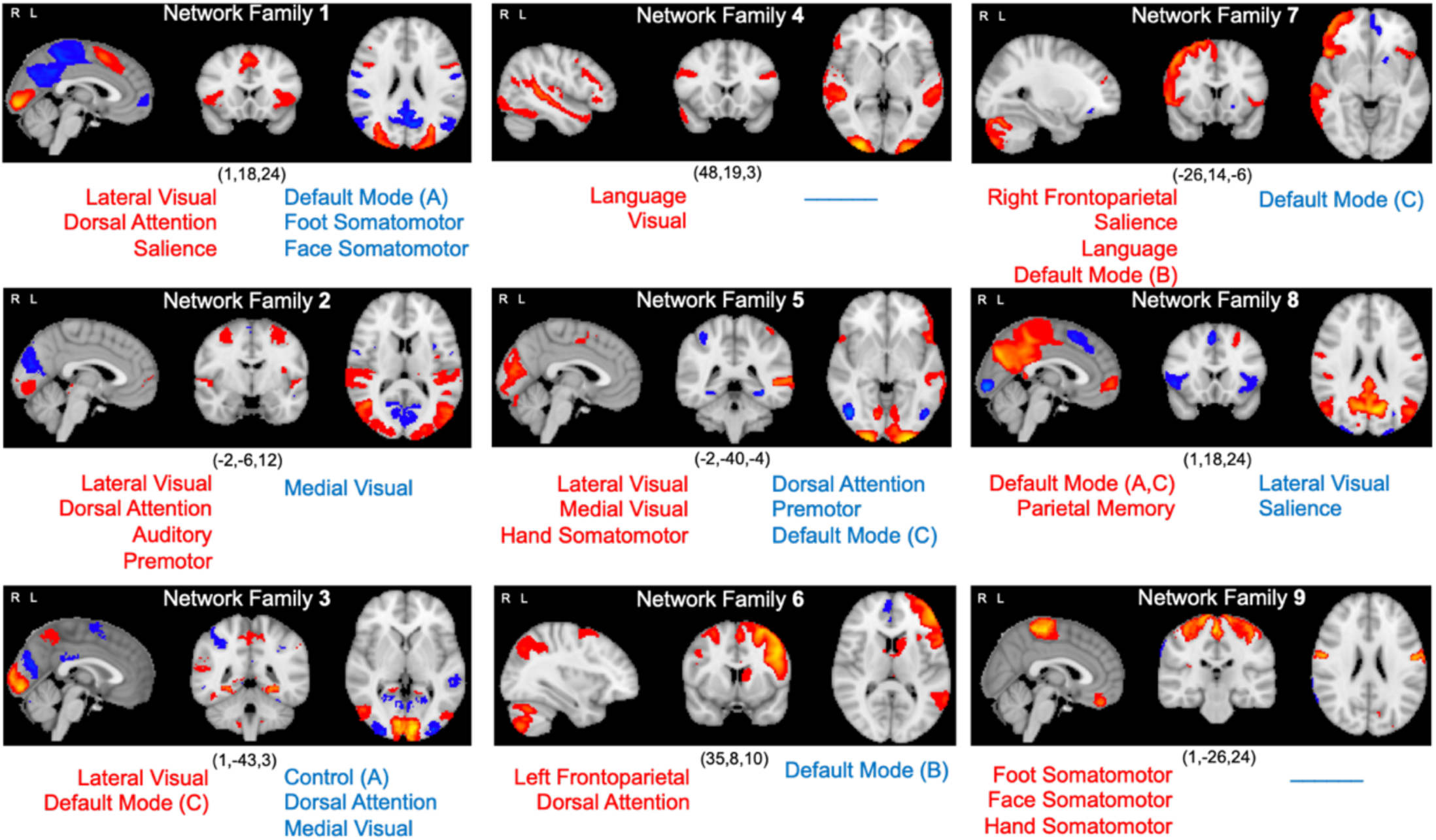
Task-evoked network family spatial maps. For each network family, the positive (red) and negative (blue) components of each network represent the average of tICA components in that family (each map thresholded at p=0.5 using a Gaussian-Gamma mixture model) to compute a mean network family spatial map. Overlap between the positive and negative components of each network family map and each of 17 canonical large-scale networks defined in the Gordon et al. parcellation^49^ was computed using the Network Correspondence Toolbox (NCT)^50^. Gordon network names listed in red showed significant overlap with positive task network components and networks listed in blue showed significant overlap with negative task network components.

### Brain Network Activity Predominantly Reflects Cognition Across All Tasks

Consistent with prior work using other fMRI-based brain features^1,4,7–24^, univariate associations between tICA network loadings and latent mental health factor scores were substantially stronger for cognition than for non-cognitive domains (negative affect, positive affect, processing speed, substance use) across all networks and all tasks (**Fig. 3**). Only cognitive associations survived correction for multiple comparisons (Bonferroni threshold at p<0.0001 to correct for 77 networks * 5 mental health domains = 385 tests) and replicated in the holdout group (Group 2); non-cognitive associations did not replicate across groups even at uncorrected thresholds. Analyses controlled for age and sex and took into account covariance in family structure within the participant sample (i.e., family members with non-independent data) using permutation analysis of linear models (PALM)^51^ with multi-level exchangeability blocks^52^. Families were not split across Group 1 and Group 2 to avoid data leakage.

**Figure 3.**
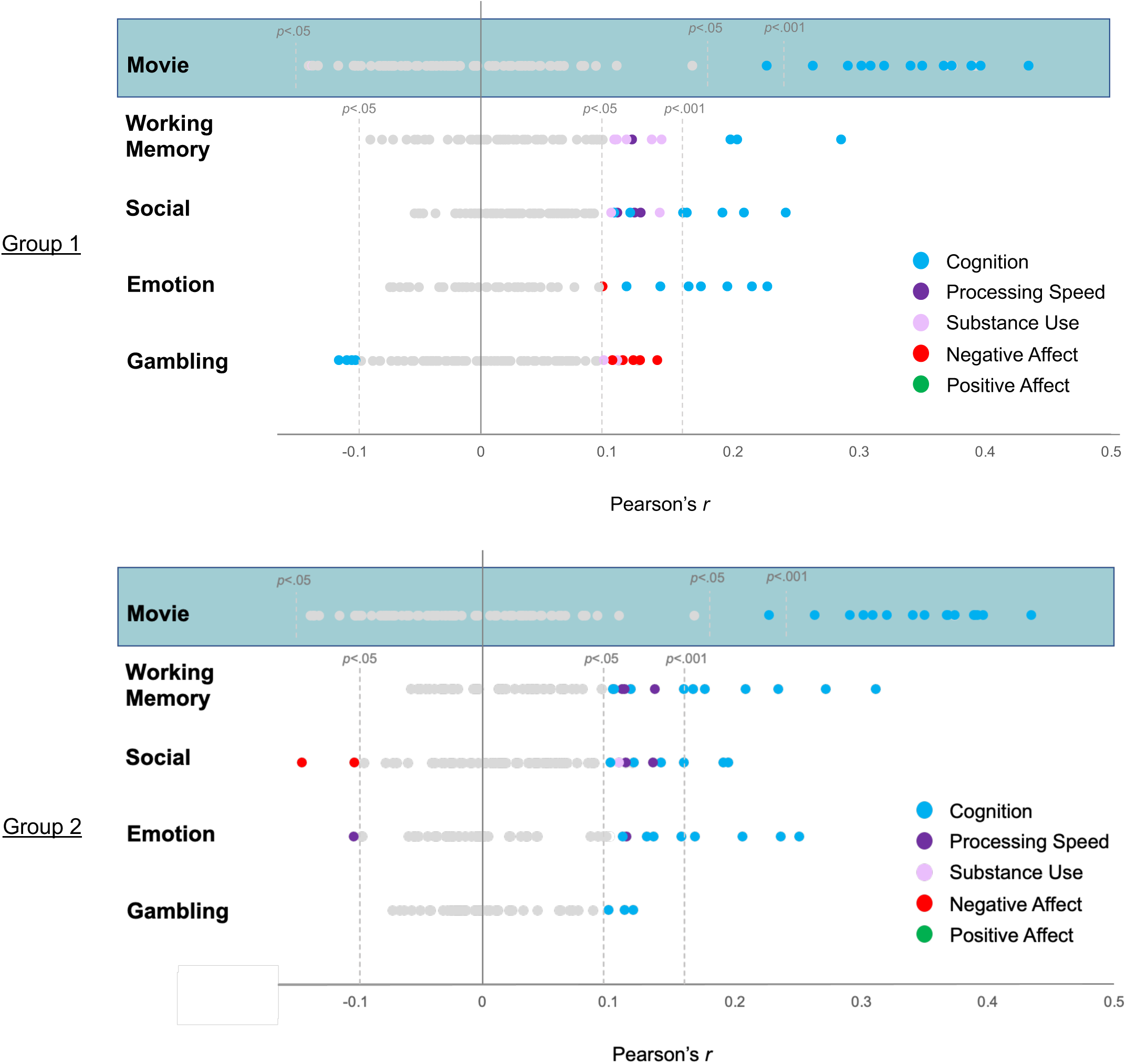
Univariate links between task-evoked network activation strengths and mental health factor scores. For each of the five tasks (rows), the Pearson’s correlation between the activation strength of each network evoked by the task and each of the five mental health factor scores is shown (dots). Significance was assessed both at the uncorrected p<.05 and p<.001 levels, and Bonferroni-adjusted p<.0001 level to correct for multiple comparisons. All relationships significant at the p<.05 level or less are color-coded by mental health domain. All p>0.05 uncorrected relationships are grayed out. Findings are presented separately for Group 1 and Group 2. Movie-watching data was obtained in only a subset of HCP-YA subject (n=184) and therefore was not split across groups. Data from this subset is visualized on both the Group 1 and Group 2 figures for easy visualization. Note that only the cognition factor yielded Bonferroni-corrected significance.

Even when considering findings at the uncorrected p<.05 or p<.001 levels, activity in five of the nine Network Families (2, 3, 5, 6 and 7) exclusively reflected cognition, regardless of task (**Fig. 1**). In Group 1, only Network Families 1, 8 and 9 reflected negative affect or substance use at the uncorrected level. However, none of these relationships replicated in Group 2. Moreover, links with positive affect (PVS) were not observed even at the uncorrected level in either Group 1 or Group 2. In both Groups 1 and 2, the Movie-Watching task elicited brain activity with the strongest link to individual differences in cognition by a wide margin, followed by the Working Memory task (**Fig. 3**). Conversely, in both groups, brain activity during the Gambling task had the weakest links with individual differences in mental health. While Gambling, Working Memory and Social Cognition task activation linked to substance use in Group 1, only brain network activity during the Social Cognition task displayed a link to individual differences in substance in both groups, albeit at a modest level (p<.05) and in different brain networks.

### Multivariate Brain Function Profiles Predict Cognition and Processing Speed but not Affect or Substance Use

Next, we examined whether considering subjects’ multivariate profile of brain activity across all tasks and evoked networks improved recovery of non-cognitive mental health signal. In order to facilitate training and testing across consistent networks in Groups 1 and 2, we restricted this analysis to the 46 Emotion, Gambling, Working Memory, and Social Cognition task networks that were identified across both groups (**Supplementary Table S1**), and to the subjects (Group 1: n=324; Group 2: n=330) who had complete data across all four tasks. Movie-watching networks were not included in this analysis given the smaller subset of subjects with this data (n=184 total). We trained a random forest machine learning regression model on Group 1 to make individualized predictions in Group 2 of each mental health factor score based on subjects’ brain function profiles. We subsequently applied SHAP (SHapely Additive exPlanations)^53^ to the predictions made by each model to quantify the feature importance of each brain network for predicting scores in each mental health domain.

In the test set (Group 2), a subject’s profile of brain activity across the 46 task-evoked networks significantly predicted their Cognition score (correlation between actual and predicted score: *r*=.15, *p*=.005, MSE=0.919) and Processing Speed score (*r*=.11, *p*=.041, MSE=1.184) to modest degrees. SHAP analysis (**Fig. 4a**) revealed that the network with the greatest feature importance for predicting Cognition was a variant from Network Family 9, evoked during the n-back Working Memory task, which involved left-lateralized motor cortex, rostral ventral putamen, and orbitofrontal cortex, as well as bilateral hippocampus and ventromedial prefrontal and ventral visual areas, negatively correlated with the right frontoparietal control network, the anterior and posterior dorsal midline and cerebellum (**Fig. 4b-c**). Higher loadings on this network were associated with better cognition. The network with the greatest feature importance for predicting processing speed was a variant from Network Family 6 (**Fig. 4a**), which involved bilateral frontoparietal and dorsal attention networks, negatively correlated with the default mode network (**Fig. 4b-c**). Multivariate brain network activity profiles did not significantly predict Positive Affect (r=.003, *p*=.956, MSE=1.255), Negative Affect (r=.03, *p*=.630, MSE=1.059) or Substance Use (r=.02, *p*=.681, MSE=1.226).

**Figure 4.**
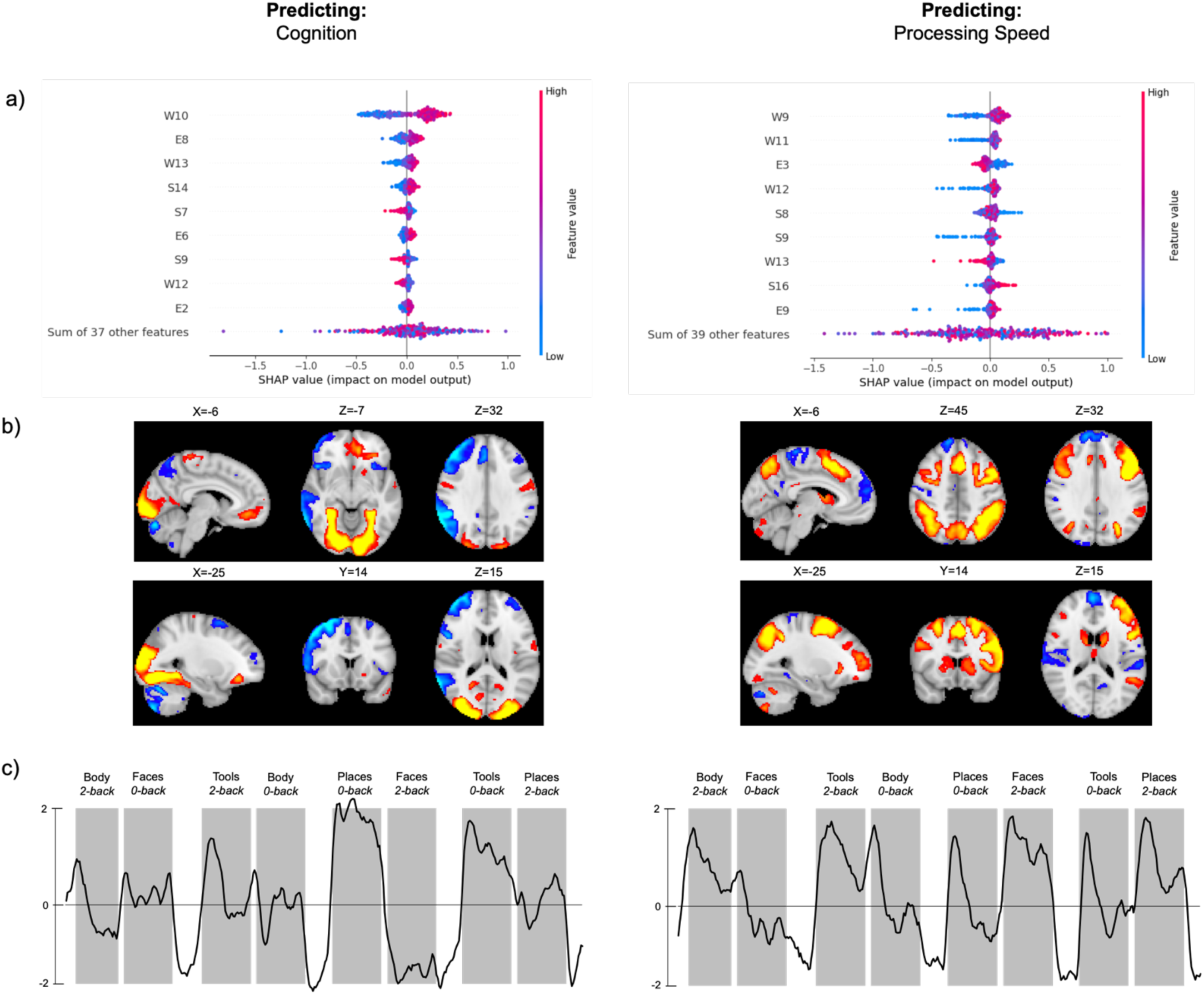
Task-evoked brain networks predicting Cognition and Processing Speed. a) Feature importance of task evoked networks for predicting Cognition (left panel) and Processing Speed (right panel). Networks are ordered from top-to-bottom on average feature importance. Each dot represents a subject’s SHAP value for a given network. Dots with positive SHAP values denote data points that increase the prediction for a given network; dots with negative SHAP values indicate data points that decrease the prediction for that network. Red/blue indicates whether the feature is positive or negative for that data point. b) Spatial map and c) time course of the network with the highest feature importance for predicting Cognition (left panel) and Processing Speed (right panel). Network time courses in c) are overlayed on the Working Memory task stimulation blocks for reference; white spaces indicate inter-block (i.e., rest) intervals. The time course in the left panel of c) (Cognition) suggests strong activation of frontoparietal areas and suppression of medial visual areas during faces 2-back vs low activation during most 0- and 2-back conditions, except for places and tools 0-back, which have positive activity of the network (e.g., strong activation of visual areas). The time course in right panel of c) (Processing Speed) suggests strong activity of the network throughout 2-back conditions and at the start of 0-back conditions.

### Evaluating the Role of Measurement Reliability

Having established the discrepancy between cognitive versus non-cognitive signal in these data, we next systematically investigated potential contributing factors. First, we examined differences in measurement reliability between cognitive and non-cognitive mental health domains. To assess this, we leveraged the HCP-YA’s Test-Retest sample (n=46) to compute the intraclass correlation (ICC) of each of the five mental health factors. ICC is the predominant index of measurement reliability in the clinical neuroimaging literature^28,54,55^, and quantifies the percent of total variance (i.e., between-subject, within-subject, and error variances) attributable to between-subject variance^56^. Retest subjects conducted their second set of HCP-YA assessments an average of 139.3 days (SD=69.0 days) after conducting their first set of assessments. The retest protocol included 78 of the 87 behavioral and neuropsychiatric assessments included in the initial testing protocol, with all nine of the missing variables belonging to the substance abuse factor. While the factor analysis procedure we employed^29^ imputes missing values, we elected not to interpret ICC values for the substance abuse factor since only two of the eleven initial substance use measurements were available in the retest data.

ICC(3,1) values for the remaining four mental health factors were uniformly high (>0.9; **Fig. 5a**), consistent with prior work showing that composite scores can yield higher reliability than their constituent measures^30,31^. These results indicate that the large difference in cognitive versus non-cognitive mental health signal observed in task-evoked brain activity is unlikely to be explained by differences in measurement reliability.

**Figure 5.**
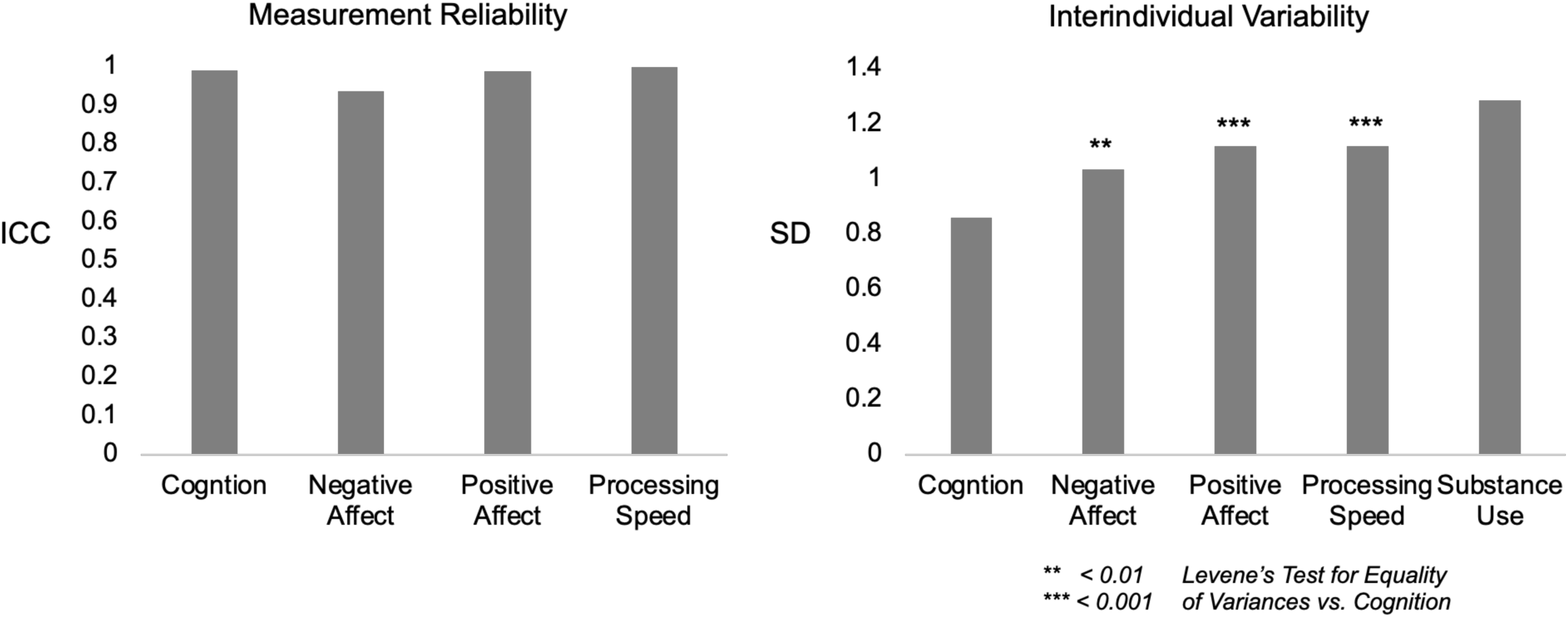
Measurement Reliability and Interindividual Variability Across Behavioral Domains. Intraclass correlation coefficients (ICC; left) and standard deviations (SD; right) for each behavioral factor. All domains demonstrated high reliability (ICC > 0.9). Cognition exhibited significantly lower interindividual variability than non-cognitive domains yet showed the strongest brain function associations, ruling out insufficient behavioral heterogeneity as an explanation for weak non-cognitive brain-behavior relationships.

### Evaluating the Role of Phenotype Dispersion

Given that HCP-YA comprises generally healthy young adults, we tested whether weak non-cognitive brain associations could be explained by restricted phenotypic heterogeneity. We found that non-cognitive mental health factor scores displayed *greater* dispersion than cognition, whether quantified by factor score standard deviation (1.2-15x larger SDs; **Fig. 5b**) or by the average coefficient of variation (CV) for individual assessment items contributing to each factor: (CV_COGNITION_=0.16; CV_PROCESSING SPEED_=0.18; CV_POSITIVE AFFECT_=0.18; CV_NEGATIVE AFFECT_=0.47; CV_SUBSTANCE USE_=2.21). This pattern replicated in Group 2. That cognition remained the most recoverable phenotype from task-evoked brain features despite its comparatively lower interindividual dispersion provides evidence against restricted phenotypic range as an appreciable contributor to weak non-cognitive mental health signal in task evoked brain activity.

### Evaluating the Role of Brain Feature Dimensionality

Next, to test whether weak non-cognitive brain–phenotype relationships may reflect insufficient representational capacity in the brain feature space, we performed a brain-dimensionality sweep (**Fig. 6a**). For each split-half group, we applied principal component analysis (PCA) to the 46-dimensional task-evoked network loading profiles and predicted each latent mental health factor score from the first *k* brain PCs (*k* = 1–46) using ridge regression with 5-fold cross-validation (CV). Increasing brain dimensionality produced domain-specific patterns. In Group 1, cognition was the only target with positive, increasing recoverability, reaching best CV *R*^2^ = 0.096 at *k* = 19. In contrast, all non-cognitive factors were at-or-below chance across the sweep and achieved their best (least negative) performance at *k* =1: Internalizing *R*^2^ = −0.006, Well-Being *R*^2^ = −0.010, Processing Speed *R*^2^ = −0.035, Substance Use *R*^2^ = −0.007. This pattern replicated in Group 2, where cognition again showed the only positive predictive signal (best CV *R*^2^ = 0.043 at *k* = 3), while non-cognitive factors remained negative and again uniformly peaked at *k* = 1: Internalizing *R*^2^ = −0.020, Well-Being *R*^2^ = −0.023, Processing Speed *R*^2^ = −0.076, Substance Use *R*^2^ = −0.011. Thus, expanding brain feature dimensionality improved recovery of cognitive but not non-cognitive mental health signal, a pattern more consistent with the brain data’s representational misalignment with non-cognitive mental health, rather than with insufficient brain feature dimensionality.

**Figure 6.**
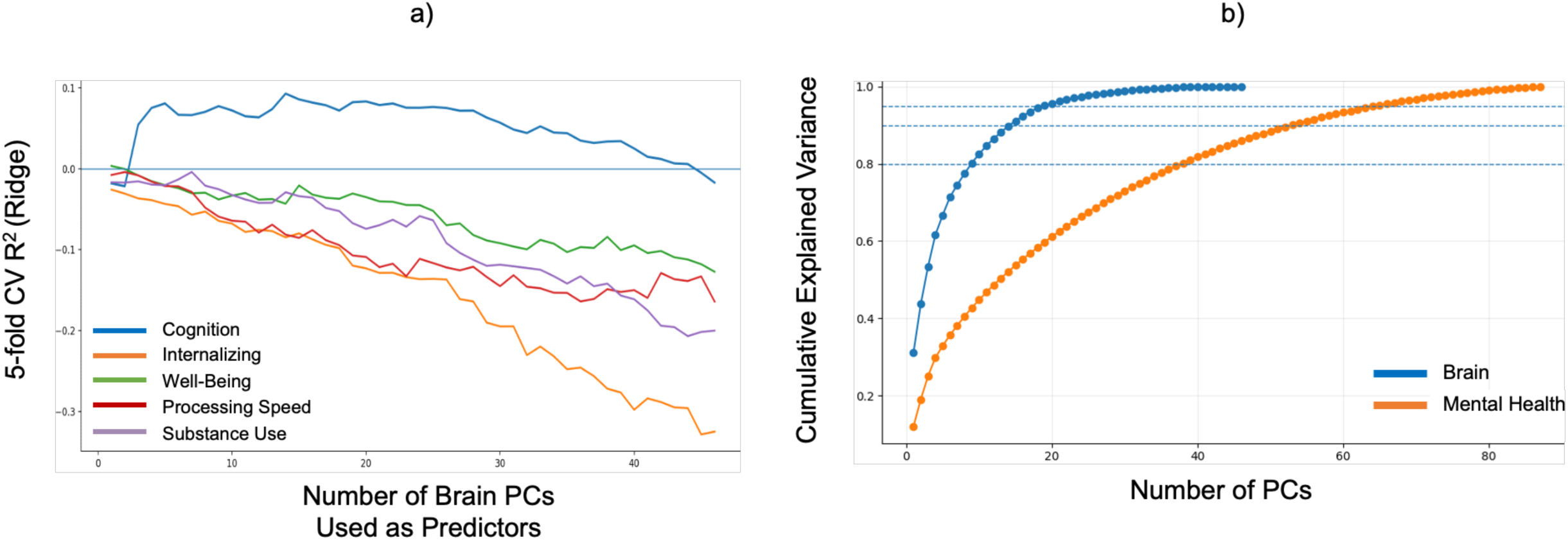
Brain space dimensionality and mental health signal. a) Increasing dimensionality of the brain feature space enhances cognitive but not non-cognitive mental health signal. b) The mental health space has higher dimensionality than the task-evoked brain activity space.

Moreover, the task-evoked brain feature space was also markedly more condensed than the mental health space: 15 PCs captured 90% of variance in the evoked brain activity profiles, whereas 53 PCs were required to capture 90% of variance in the mental health profiles (**Fig. 6b**).

### Comparing Interindividual Variability in Evoked Brain Signal and Mental Health

We next quantified how much between-subject heterogeneity is expressed in the task-evoked brain feature space versus the behavioral phenotyping space by computing pairwise similarity among subjects’ multivariate profiles. We calculated subject–subject profile correlations within each space and summarized overall homogeneity using the mean absolute correlation and mean shared variance (mean *r*^2^) across all subject pairs. Subjects were substantially more similar to one another in brain space than in mental health space, t(101779)=109.64, *p*<.001. In Group 1, the mean absolute subject–subject correlation was 0.224 for brain profiles versus 0.132 for behavioral profiles (1.70×higher), and mean shared variance was 0.075 versus 0.027 (2.75× higher). In Group 2, this pattern replicated with mean absolute correlations of 0.216 for brain profiles versus 0.136 for behavioral profiles (1.59× higher) and mean shared variance of 0.070 versus 0.029 (2.42× higher). To rule out an artifact of comparing 46 brain features to 87 behavioral measures, we repeated the subject–subject similarity analysis after randomly subsampling 46 behavioral variables (1000 draws). Task-evoked brain profiles still showed 2.0× higher mean pairwise shared variance (mean *r*^2^) and 1.45× higher mean absolute profile similarity than dimensionality-matched behavioral profiles, indicating the effect is not predominantly driven by feature count. Finally, separating out cognitive and non-cognitive mental health domains showed they had similarly low levels of between-subject similarity compared to the brain data (cognitive: r=0.174; non-cognitive: r=0.161. Thus, compared to the mental health space, the task-evoked brain space is more homogeneous/redundant across individuals, consistent with reduced overall subject discriminability and a constrained individual-difference representation in the task-evoked features.

### Brain Function Subtypes and Mental Health Subtypes Have Poor Correspondence

Finally, we examined the consequences of misalignment between task evoked brain activity and mental health for biotyping concordance and informativeness. To do so, we applied hierarchical clustering, separately, to subjects’ brain function profiles and mental health profiles (**Fig. S1**). This clustered each individual into one subtype based on their brain function profile and one subtype based on their mental health profile. Then, we assessed the correspondence between these two sets of subtypes at all levels where the number of clusters was common to both the brain function and mental health subtyping sets (i.e., k=2, 5 and 10; **Fig. S1**). The mental health characteristics of each subtyping solution are presented in **Supplementary Figure S2**.

Brain function subtypes and mental health subtypes were largely non-corresponding: a given subject’s brain function subtype was not predictive of their mental health subtype, and vice-versa (**Table 1**, **Fig. 7**). While subtype correspondence was statistically significant at the k=2 clustering level (*Χ*²(1)=4.49, *p*=.034), Table 1 illustrates the low effect size and specificity of the overlap. For instance, of 218 subjects in brain function subtype 1, 42% were in mental health subtype 1 and 58% were in mental health subtype 2. Correspondence did not differ from chance at finer clustering levels (k=5: *Χ*²(16)=23.81, *p*=.094; k=10: *Χ*²(16)=76.91, *p*=.608).

**Figure 7.**
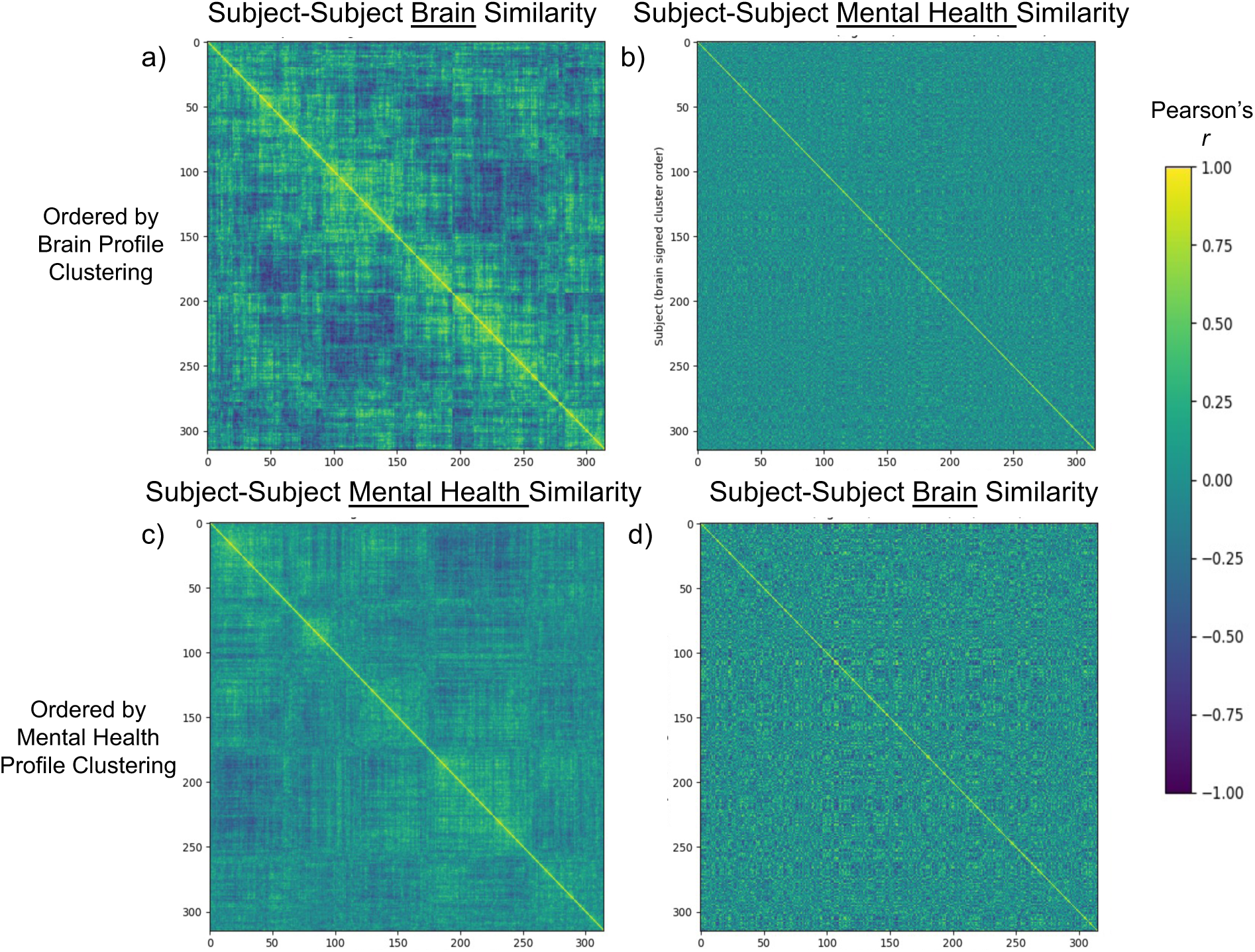
Non-alignment of brain function subtypes and mental health subtypes. Subjects with similar evoked brain activity profiles (a) had little similarity in mental health profile (b), and vice versa: subjects with similar mental health profiles (c) had little similarity in evoked brain activity profile (d).

**Table 1.**
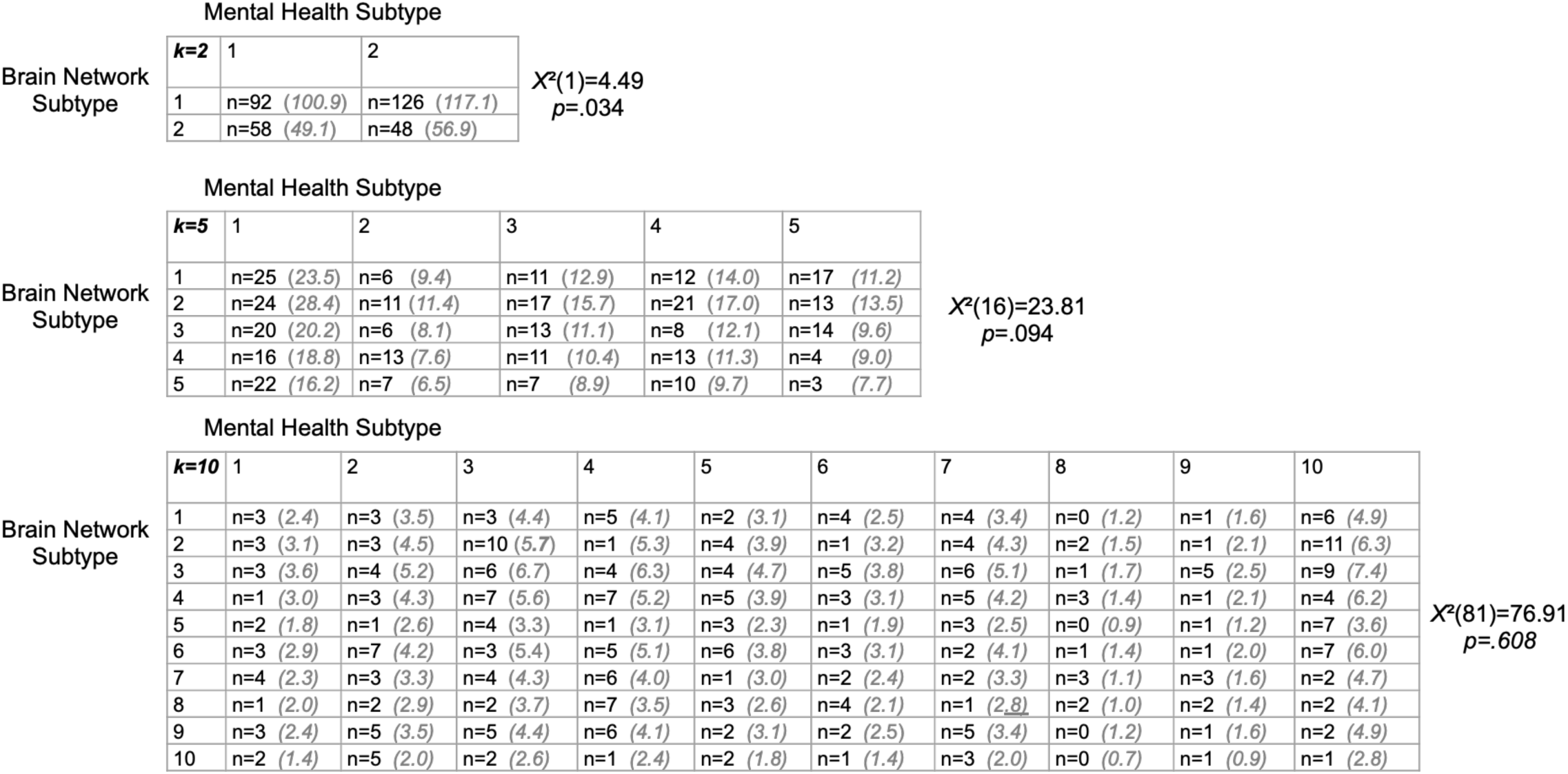
Non-alignment of brain function subtypes and mental health subtypes. At each clustering solution level common to both the brain function and mental health subtyping sets (k=2, 5 and 10), the total count (n) of subjects with each joint brain function subtype–mental health subtype assignment. Grayed values indicate the expected count if joint subtype assignments were random. Chi-squared tests examining the difference between the expected (by random chance) and actual joint assignment counts indicated that subjects in a brain function subtype were randomly distributed across mental health subtypes, and vice versa, underscoring the non-alignment of the subtypes.

|  |  | Mental Health Subtype |  |  |  |
| --- | --- | --- | --- | --- | --- |
| Brain Network Subtype | $k=2$ | 1 | 2 | | |
| | 1 | n=92 (100.9) | n=126 (117.1) | $X^2(1)=4.49$<br>$p=.034$ | |
|  | 2 | n=58 (49.1) | n=48 (56.9) |  |  |

|  |  | Mental Health Subtype |  |  |  |  |  |  |
| --- | --- | --- | --- | --- | --- | --- | --- | --- |
| Brain Network Subtype | $k=5$ | 1 | 2 | 3 | 4 | 5 | $X^2(16)=23.81$<br>$p=.094$ | |
|  | 1 | n=25 (23.5) | n=6 (9.4) | n=11 (12.9) | n=12 (14.0) | n=17 (11.2) |  |  |
|  | 2 | n=24 (28.4) | n=11 (11.4) | n=17 (15.7) | n=21 (17.0) | n=13 (13.5) |  |  |
|  | 3 | n=20 (20.2) | n=6 (8.1) | n=13 (11.1) | n=8 (12.1) | n=14 (9.6) |  |  |
|  | 4 | n=16 (18.8) | n=13 (7.6) | n=11 (10.4) | n=13 (11.3) | n=4 (9.0) |  |  |
|  | 5 | n=22 (16.2) | n=7 (6.5) | n=7 (8.9) | n=10 (9.7) | n=3 (7.7) |  |  |

**Table 1.** *Non-alignment of brain function subtypes and mental health subtypes.*
|  |  | Mental Health Subtype |  |  |  |  |  |  |  |  |  |  |  |
| --- | --- | --- | --- | --- | --- | --- | --- | --- | --- | --- | --- | --- | --- |
| Brain Network Subtype | $k=10$ | 1 | 2 | 3 | 4 | 5 | 6 | 7 | 8 | 9 | 10 | $X^2(81)=76.91$<br>$p=.608$ | |
|  | 1 | n=3 (2.4) | n=3 (3.5) | n=3 (4.4) | n=5 (4.1) | n=2 (3.1) | n=4 (2.5) | n=4 (3.4) | n=0 (1.2) | n=1 (1.6) | n=6 (4.9) |  |  |
|  | 2 | n=3 (3.1) | n=3 (4.5) | n=10 (5.7) | n=1 (5.3) | n=4 (3.9) | n=1 (3.2) | n=4 (4.3) | n=2 (1.5) | n=1 (2.1) | n=11 (6.3) |  |  |
|  | 3 | n=3 (3.6) | n=4 (5.2) | n=6 (6.7) | n=4 (6.3) | n=4 (4.7) | n=5 (3.8) | n=6 (5.1) | n=1 (1.7) | n=5 (2.5) | n=9 (7.4) |  |  |
|  | 4 | n=1 (3.0) | n=3 (4.3) | n=7 (5.6) | n=7 (5.2) | n=5 (3.9) | n=3 (3.1) | n=5 (4.2) | n=3 (1.4) | n=1 (2.1) | n=4 (6.2) |  |  |
|  | 5 | n=2 (1.8) | n=1 (2.6) | n=4 (3.3) | n=1 (3.1) | n=3 (2.3) | n=1 (1.9) | n=3 (2.5) | n=0 (0.9) | n=1 (1.2) | n=7 (3.6) |  |  |
|  | 6 | n=3 (2.9) | n=7 (4.2) | n=3 (5.4) | n=5 (5.1) | n=6 (3.8) | n=3 (3.1) | n=2 (4.1) | n=1 (1.4) | n=1 (2.0) | n=7 (6.0) |  |  |
|  | 7 | n=4 (2.3) | n=3 (3.3) | n=4 (4.3) | n=6 (4.0) | n=1 (3.0) | n=2 (2.4) | n=2 (3.3) | n=3 (1.1) | n=3 (1.6) | n=2 (4.7) |  |  |
|  | 8 | n=1 (2.0) | n=2 (2.9) | n=2 (3.7) | n=7 (3.5) | n=3 (2.6) | n=4 (2.1) | n=1 (2.8) | n=2 (1.0) | n=2 (1.4) | n=2 (4.1) |  |  |
|  | 9 | n=3 (2.4) | n=5 (3.5) | n=5 (4.4) | n=6 (4.1) | n=2 (3.1) | n=2 (2.5) | n=5 (3.4) | n=0 (1.2) | n=1 (1.6) | n=2 (4.9) |  |  |
|  | 10 | n=2 (1.4) | n=5 (2.0) | n=2 (2.6) | n=1 (2.4) | n=2 (1.8) | n=1 (1.4) | n=3 (2.0) | n=0 (0.7) | n=1 (0.9) | n=1 (2.8) |  |  |

### Subtyping Misalignment Yields Groups with Discordant Brain Function-Mental Health Associations

We next examined the consequence of this subtyping misalignment for group-level analyses of brain function-mental health associations. We found that the starting point of analysis (i.e., whether to subtype subjects by mental health data or by brain activation data) meaningfully altered conclusions about brain function-mental health group-level associations, in sometimes opposite directions (**Fig. 8**).

**Figure 8.**
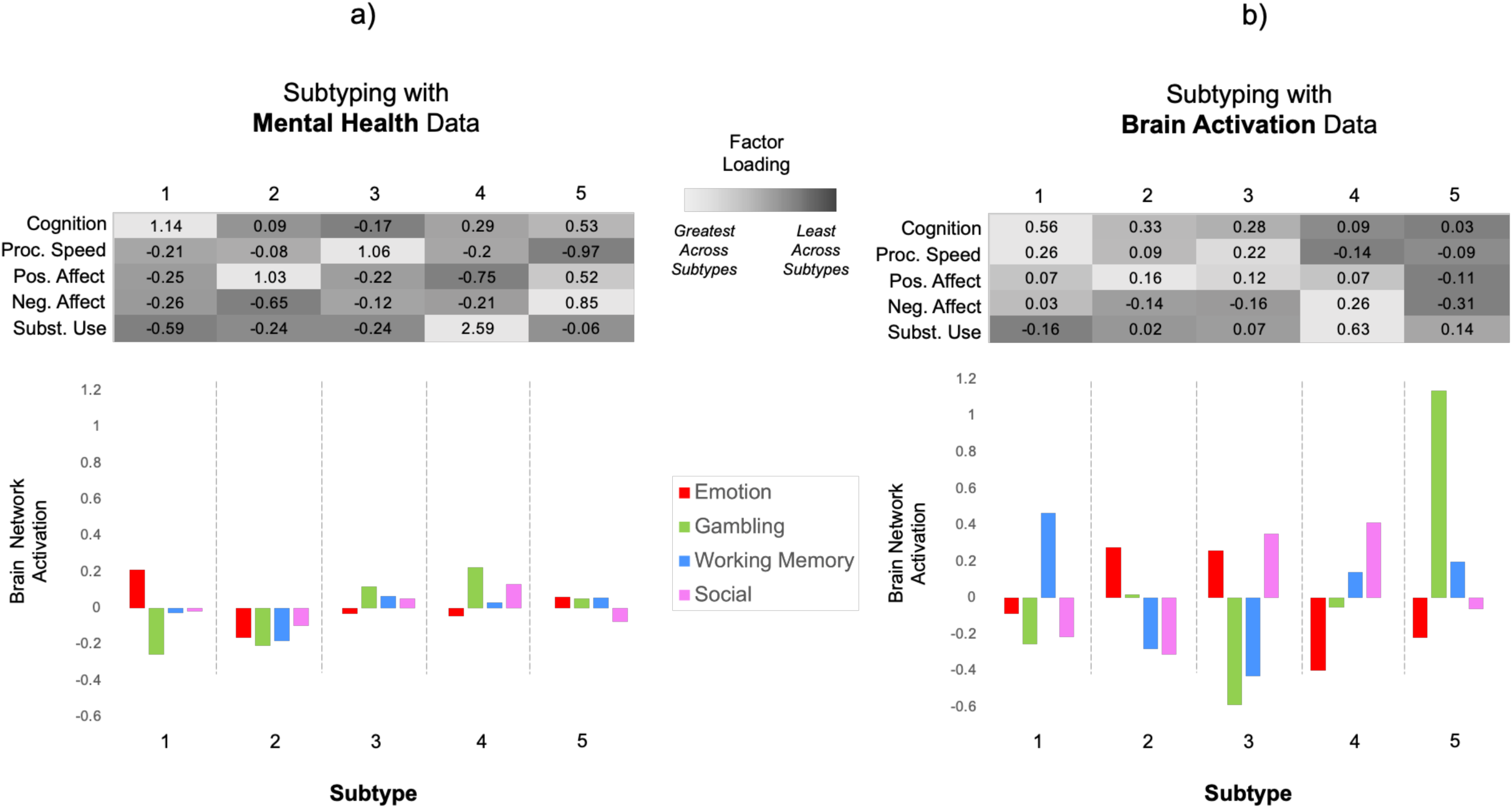
Comparing brain function-mental health associations across subtyping solutions. Average mental health profile (top panel; *Z*-scores) and average brain function profile (bottom panel; *Z*-scores) of each subtype derived from clustering of a) mental health data and b) brain function data at the k=5 clustering solution level. Given that within-subject loadings on networks activated by the same task were strongly correlated, such that participants with high loadings on one pattern generally showed high loadings on other patterns (**Fig. S3**), for this analysis we operationalized brain network activation on each task as the *z*-scored mean subject loading across all networks activated by the task.

For instance, in the k=5 mental health subtyping set (**Fig. 8a**), the high cognition, low substance use group (subtype 1) was characterized at the brain level by high network loadings during the Emotion task and low network loadings during the Gambling task, with average network loadings on the Working Memory and Social Cognition tasks (**Fig. 8a**). But in the k=5 brain function subtyping set (**Fig. 8b**), the analogous high cognition, low substance use group (subtype 1) was characterized at the brain level by very high network loadings during the Working Memory task and average network loadings during the Emotion task. Similarly, in the mental health subtyping set, the high substance use group (subtype 4) was characterized at the brain level by high network loadings during the Gambling task and average network loadings during the Emotion task, whereas in the brain function subtype set, the high substance use group (subtype 4) was characterized at the brain level by average network loadings during the Gambling task and very low network loadings during the Emotion task. Overall, due to the discordance of the mental health and brain function data, the starting point of analysis (i.e., whether to subtype subjects by mental health or brain function) meaningfully alters group-level associations between brain function and mental health.

### Brain Function Subtypes Delineate Mental Health Differences Better than Mental Health Subtypes Delineate Brain Function Differences

Finally, given the observed discordance between subtyping sets, we compared the relative ability of subtypes generated by one modality to yield groups that significantly differentiated facets of the other. Independent samples t-tests at the k=2 clustering level (controlling for age and sex, and accounting for covariance in family structure using PALM) demonstrated that subtypes generated using brain function data had significantly different means of cognition (*t*(322)=3.18, p<.001), processing speed (*t*(322)=2.27, p=.014), and substance use (*t*(322)=2.77, p=.002), but not of negative affect/internalizing (*t*(322)=0.71, p=.790) or positive affect/well-being (*t*(322)=1.06, p=.220). On the other hand, subtypes generated using mental health data only significantly differed in brain activation on the Gambling task (*t*(322)=2.84, p=.002), and not in brain activation on the Emotion (*t*(322)=0.72, p=.258), Working Memory (*t*(322)=1.13, p=.110) or Social Cognition (*t*(322)=1.56, p=.064) tasks. Similarly, one-way ANOVAs at the k=5 clustering level demonstrated that subtypes generated using brain function data had significantly different means of cognition (F(6,316)=3.84, p=.004), internalizing (F(6,316)=2.67, p=.030), and substance use (F(6,316)=3.20, p=.014), but not of positive affect (F(6,316)=0.53, p=.733) or processing speed (F(6,316)=1.49, p=.240). On the other hand, subtypes generated using mental health data only significantly differed in brain activation on the Gambling task (F(6,316)=3.03, p=.016), and not in brain activation during the Emotion (F(6,316)=2.02, p=.110), Working Memory (F(6,316)=1.16, p=.328) or Social Cognition (F(6,316)=0.77, p=.565) tasks. Overall, these findings indicate brain function data were more sensitive to mental health differences than mental health data were sensitive to brain function differences.

## Discussion

The low reliability of early fMRI findings pushed the neuroimaging field toward the implementation of simple tasks in large datasets to enhance the reproducibility of brain activations. Yet, Hedge and colleagues describe a “reliability paradox”^38^ built into these fMRI tasks, in which tasks designed to elicit robust group-level brain activations to map task-relevant brain regions inherently have low between-subject variability in brain activity – which places a constraint on individual differences research. Still, we demonstrate here that the magnitude of between-subject variability in task fMRI-evoked brain activity in generally healthy adults can be sufficient – with large enough sample sizes and optimized methodology – to robustly parse individual differences even in low-variability target phenotypes (i.e., Cognition). What, then, could explain why the same fMRI data do not parse individual differences in other phenotypes (i.e., positive and negative affect) with comparable measurement reliability and even larger interindividual variability? We posit that it is because prevalent fMRI tasks used in large-scale neuroimaging studies, simplified to ease data collection in large populations, are not challenging or evocative enough to elicit individual differences in brain activity mapping the ecological range of affective function in healthy adults. If this is the case, continuing methodological advances in brain signal isolation, modelling optimization and spatiotemporal resolution may not meaningfully strengthen our ability to extract this information. Rather, the current findings point to a need to leverage different types of tasks or paradigm designs in future large-scale neuroimaging studies if our goal is to evoke brain activity that more holistically represents mental health.

Cardiac stress tests provide an apt analogy. Holistic assessments of cardiovascular health require not only measures of heart function at the resting or low-level activity state, but at the vigorous activity state as well. We posit that the fMRI tasks commonly adopted for large-scale studies do not take the brain particularly far away from its resting-state, and that more emotionally demanding tasks that facilitate measurement of brain function during higher arousal states will need to be more broadly implemented and incorporated into large-scale studies. fMRI tasks implemented in large-scale neuroimaging studies tend to be selected for brevity, ease of administration, and established consistency of results. This skews paradigm selection towards block designs that provide maximized signal-to-noise ratios and require less individualized analytic processing pipelines, but which often mask important nuances in brain reactivity to specific stimuli (e.g., correct versus incorrect trials) and average out significant individual differences in brain activation. While more challenging to administer consistently in a large-scale study, tasks that model subject responses and brain activity at higher detail (i.e., event-related designs) and that are more affectively provocative and challenging (e.g., social exclusion task, cold pressor task, monetary incentive delay task) may prove more powerful in extracting individual differences in non-cognitive mental health with enhanced power of larger sample sizes. Individual-differences studies using symptom-proximal manipulations—particularly psychosocial stress and social-threat/ostracism paradigms—have reported stronger associations with anxiety/depression-related traits than many canonical localizer-style tasks, suggesting that task design can meaningfully influence the detectability of non-cognitive mental health signal^57–60^. Importantly, though, paradigms involving highly salient real-life consequences must be carefully considered in the context of research ethics and subject beneficence.

Many basic science investigations, clinical trials, grant proposals and large-scale study designs rely on explicit or implicit notions that particular fMRI tasks differentially probe individual differences in distinct RDoC domains. For instance, the HCP-YA Emotion task and related variants are used specifically as probes of NVS in the Connectomes Related to Human Disease (CRHD) investigating anxiety and depression^61^, the UK Biobank^62^, and the Transdiagnostic Connectome Project (TCP)^63^; the Gambling task and related variants are commonly used as probes of PVS, as in the CRHD related to anxiety and depression; and the Working Memory task and related variants are commonly used as probes of CS, for example, in the CRHD brain aging and dementia study^64^. These tasks are indeed validated “functional localizers”^65^ of the neural substrates relevant to these domains (e.g., amygdala activation in the Emotion task). However, as the present findings underscore, just because a task activates neural substrates underlying a domain of function, does not necessarily mean that it elicits interindividual variability in the activity of these substrates that reflects variability in that domain of function. This likely reflects the accumulating evidence that brain regions are multifunctional and have roles in diverse processing streams depending on task needs. For instance, while the Emotion task robustly activated the amygdala – a structure with known roles in NVS – neither variability in networks involving the amygdala, nor amygdala connectivity or activity itself indexed interindividual variability in negative affect/internalizing^25^. Instead, it appears that individual variability in amygdala activity on this task primarily reflects variation in the amygdala’s roles in cognition. In this way, despite these tasks’ distinct names, stimuli and instructions, individual differences in brain activity evoked by the Emotion, Gambling, Working Memory, Social Cognition, and Movie-Watching paradigms all predominantly reflected variability in cognition. Therefore, these tasks may all be more appropriately categorized as paradigms in the CS domain of RDoC – at least when administered to generally healthy young adults that may have only sub-clinical psychopathology. For this population, these tasks may be similarly unchallenging, unprovocative, and easily navigable with minimal recruitment of non-cognitive neural resources. In addition, some of the tasks may map to general cognition rather than a specific cognitive domain (such as working memory). Certainly, these tasks’ ability to elicit brain activity reflecting cognition is valuable, given that cognitive deficits represent a core feature of numerous forms of mental illness. However, most disorders also involve a broader constellation of affective and other deficits. Eliciting and identifying brain activity that reflects these non-cognitive domains will remain a critical task of clinical neuroimaging moving forward. Moreover, because the RDoC framework explicitly conceptualizes psychopathology as deviations from normative function, the capacity of fMRI tasks to elicit mental health-relevant individual differences across the normative range is critical.

It remains possible that variability in non-cognitive mental health domains may be better detected in brain activity variability modelled at other scales of brain organization than large-scale networks. However, recent work demonstrated that region-of-interest activation and edge-level connectivity were no better than large-scale network activation in the Emotion task in representing non-cognitive mental health domains^25^. Moreover, our present findings show that varying the dimensionality of the brain feature space did not help to recover non-cognitive mental health signal. Collectively, we posit that the data implicates as the key contributing factor a fundamentally poor ability of fMRI tasks currently included in many large-scale studies to elicit brain activity that reflects individual differences in non-cognitive mental health domains.

Our parallel subtyping findings point to another important set of issues. Many studies begin with mental health- or behaviour-based subject groupings (e.g., a clinical sample and a control sample) and seek to identify brain differences between the groups. Yet, we show here that groups of healthy individuals subtyped by mental health profile have largely similar brain function profiles during these fMRI tasks. This converges with findings from the clinical realm wherein patients classified to distinct DSM diagnostic categories often have minimal neurobiological differentiation from one another^66^. On the other hand, subtyping these same individuals by brain function profile yielded modest discrimination of mental health profiles. For instance, brain function subtypes were able to separate out subjects with distinct levels of cognition, internalizing and substance use. In alignment with the RDoC framework, which reconceptualizes mental health as a functional continuum across multiple domains, our findings support the movement toward a neurobiologically-based clinical taxonomy instead^67^. These findings indicate that beginning analyses with brain-based subtypes rather than mental health/behaviour-based subtypes may provide a better means of identifying fundamentally similar individuals and may be more successful at targeting biologically based treatment approaches to the individuals who will best respond to them.

A potential limitation is the present study’s use of tICA to quantify brain function phenotypes. While tICA is a purely data-driven technique, its use involves some subjective decision points as described in the Methods, including the choice of model order and the classification of noise and signal components. The present study is also limited in examining generally healthy young adults, constraining the generalization of findings to clinical populations. Though, it should be noted that HCP-YA contains several hundred subjects with alcohol and cannabis use disorders, and subjects with subclinical depression and anxiety, and we found that HCP-YA subjects are more heterogeneous in negative affect/internalizing and positive affect/well-being than they are in cognition, thus making HCP-YA a sensible dataset for mapping PVS, NVS and CS across the normative range. An important aim of future work will be to explore whether the examined fMRI paradigms, and those similar to them, are more effective at eliciting brain activity reflecting non-cognitive mental health in clinical cohorts, although some studies have already shown poor performance for this purpose^68^. However, the present findings and other published research suggest that even if they are more effective in clinical cohorts, dimensional RDoC-style analyses are not possible and interpreting case-control differences using these tasks will be confounded by the failure to elicit relevant mental health signals in the normative cohort rather than a brain difference between patients and healthy controls per se. As such, given that RDoC casts mental illness as deviations from normative biobehavioral functioning, it is important to use tasks that map mental health-related variability in normative cohorts like HCP-YA to capture the full range of function. If the task batteries in these normative large-scale studies are ineffective at eliciting mental health-relevant brain activity, this creates a challenge for identifying the deviations that define mental illness using these datasets.

## Methods

### Subjects

Data used in these analyses include fMRI scans from individuals collected as part of the Human Connectome Project (HCP) Young Adult 1200 data release. A detailed description of the recruitment for the HCP is provided by others^69,70^. Briefly, individuals were excluded from participation if they reported a history of major psychiatric disorder, neurological disorder, or medical disorder known to influence brain function. For reproducibility assessment, we randomly split eligible subjects into two groups (n1=607; n2=599) with non-overlapping family structures. Given the oversampling of siblings and twins in the HCP-YA cohort, consolidating subjects from the same family within the same group prevented family structure-related data leakage across the two groups. To maximize the generalizability of replication analyses, we did not rebalance the family-separated groups by other demographic variables.

Subjects had variable data availability and quality across the different fMRI tasks examined in the present study. To maximize the quantity of analyzable data for detection of task-evoked brain networks and brain function-mental health univariate relationships, we first established task-specific groups by excluding only subjects from each task group who had missing data, HCP quality assessment flags, or excessive head motion (i.e., 1 or more frames with framewise displacement > 2mm) for that task. This produced the following sample sizes for each set of task groups: Emotion (n1=413; n2=416), Gambling (n1=444; n2=444), Social Cognition (n1=406; n2=435), Working Memory (n1=404; n2=395). Then, for multi-network predictive modeling across groups, we winnowed the task-specific groups into a set of task-general groups (n1=324; n2=330) that comprised subjects with available and high-quality data across all four tasks.

Separately, we also examined the subset of subjects (n=184) who had 7T movie-watching data. Given the smaller sample size of subjects with movie-watching data, we did not split this sample or include movie-watching data in multi-network predictive modeling for reproducibility analysis.

### fMRI Acquisition

HCP neuroimaging data were acquired with a standard 32-channel head coil on a Siemens 3T Skyra modified to achieve a maximum gradient strength of 100 mT/m^69^. T1-weighted high resolution structural images, acquired using a 3D MPRAGE sequence with 0.7mm isotropic resolution (FOV=224mm, matrix=320, 256 sagittal slices, TR=2400ms, TE=2.14ms, TI=1000ms, FA=8°), were preprocessed via the HCP minimal pre-processing pipelines to register functional MRI data to a standard brain space. Blood oxygenation level dependent (BOLD) functional magnetic resonance imaging (fMRI) data were acquired using a gradient-echo EPI sequence with the following parameters: TR = 720 ms, TE = 33.1 ms, flip angle = 52°, FOV = 280 × 180 mm, Matrix = 140 × 90, Echo spacing = 0.58 ms, BW = 2290 Hz/Px, 72 slices, 2.0 mm isotropic voxels, with a multiband acceleration factor of 8. HCP neuroimaging data were acquired in two separate imaging sessions (each with two scans in which the direction of phase encoding alternated between anterior-to-posterior and posterior-to-anterior) over a two-day study visit.

HCP movie-watching fMRI data were acquired during a separate, third session on a 7T Siemens Magnetom scanner at the Center for Magnetic Resonance Research at the University of Minnesota. Data were collected using a gradient-echo-planar imaging (EPI) sequence with the following parameters: repetition time (TR) = 1000 ms, echo time (TE) = 22.2 ms, flip angle = 45 deg, field of view (FOV) = 208 × 208 mm, matrix = 130 × 130, spatial resolution = 1.6 mm^3^, number of slices = 85, multiband factor = 5, image acceleration factor (iPAT) = 2, partial Fourier sampling = 7/8, echo spacing = 0.64 ms, bandwidth = 1924 Hz/Px. The direction of phase encoding alternated between anterior-to-posterior (AP: MOVIE1, MOVIE4) and posterior-to-anterior (PA: MOVIE2, MOVIE3).

### fMRI Tasks

Detailed descriptions of the HCP-YA fMRI task paradigms can be found here^7,71^. Brief summaries of each task paradigm are provided below.

The Emotion task (i.e., emotional face-matching task) was adapted from Hariri and colleagues^72^. Subjects were presented with blocks of trials that asked them to decide either which of two faces presented on the bottom of the screen matched the face at the top of the screen, or which of two shapes presented at the bottom of the screen matched the shape at the top of the screen. The faces had either angry or fearful expressions. Each run included blocks of 6 trials of the same task (face or shape), with the stimulus presented for 2 s and a 1 s ITI. Each block was preceded by a 3 s task cue (“shape” or “face”), so that each block was 21 s including the cue. Each of the two runs included 3 face blocks and 3 shape blocks.

The Gambling task (i.e., incentive processing task) was adapted from Delgado and Fiez^73^. Subjects guessed the number on a mystery card in order to win or lose money. Card numbers ranged from 1–9, and they were told to indicate if they thought the mystery card number was more or less than 5 by pressing one of two buttons on the response box. Feedback consisted of the number on the card (generated by the program as a function of whether the trial was a reward, loss or neutral trial) and either: 1) a green up arrow with “$1” for reward trials, 2) a red down arrow next to −$0.50 for loss trials; or 3) the number 5 and a gray double headed arrow for neutral trials. The task was presented in blocks of 8 trials that were either mostly reward (6 reward trials pseudo randomly interleaved with either 1 neutral and 1 loss trial, 2 neutral trials, or 2 loss trials) or mostly loss (6 loss trials interleaved with either 1 neutral and 1 reward trial, 2 neutral trials, or 2 reward trials). All participants were provided with money for completing the task, though it was a standard amount across subjects.

The Social Cognition task (i.e., theory of mind task) was adapted from Castelli and colleagues^74^ and Martin and colleagues^75^. Subjects watched short video clips (20 s) of objects (squares, circles, triangles) either interacting in some way, or moving randomly. After each video clip, subjects chose between 3 possibilities: whether the objects had a social interaction (an interaction that appeared as if the shapes were considering each other’s feelings and thoughts), Not Sure, or No interaction (i.e., there was no obvious interaction between the shapes and the movement appeared random). The task included 5 video blocks (2 Mental and 3 Random in one run, 3 Mental and 2 Random in the other run) and 5 fixation blocks (15 s each).

The Working Memory task (i.e., N-back task) involved blocks of trials that consisted of pictures of faces, places, tools and body parts. Within each run, the 4 different stimulus types were presented in separate blocks within the run. Within each run, ½ of the blocks consisted of a 2-back working memory task (respond ‘target’ whenever the current stimulus is the same as the one two back) and ½ presented a 0-back working memory task (a target cue is presented at the start of each block, and the person must respond ‘target’ to any presentation of that stimulus during the block). A 2.5 s cue indicated the task type (and target for 0-back) at the start of the block. Each of the two runs contains 8 task blocks (10 trials of 2.5 s each, for 25 s) and 4 fixation blocks (15 s each). On each trial, the stimulus was presented for 2 s, followed by a 500 ms ITI. Each block contained 10 trials, of which 2 were targets, and 2–3 were non-target lures (e.g., repeated items in the wrong n-back position, either 1-back or 3-back). The inclusion of lures was critical to ensure that participants were using an active memory approach to the task and allowed one to assess conflict related activity as well as error related activity.

The Movie-Watching task involved passively viewing a series of video clips with audiovisual content. Each MOVIE run consisted of 4 or 5 clips, separated by 20 s of rest (indicated by the word “REST” in white text on a black background). The MOVIE1 run examined here contained clips from independent films (both fiction and documentary) made freely available under Creative Commons license on Vimeo, as well as a final clip comprising a montage of brief (1.5 s) videos that was identical across each of the four Movie-Watching runs (to facilitate test-retest and/or validation analyses). We focused on MOVIE1 because it was the only Movie-Watching run previously found to elicit functional brain organization predictive of both cognition and emotion^7^. Audio was delivered via Sensimetric earbuds.

### fMRI Preprocessing

For the Emotion, Gambling, Social Cognition and Working Memory task fMRI data, we began with the HCP’s minimally preprocessed data. The HCP minimal preprocessing pipeline for fMRI data removes spatial distortions, realigns volumes to compensate for subject motion, registers the echo planar functional data to the structural data, reduces the bias field, normalizes the 4D image to a global mean, and masks the data with a final FreeSurfer-generated brain mask^76^. We additionally applied 4mm spatial smoothing, 150 sec temporal smoothing, removal of non-steady-state frames (first 15 frames), and ICA-FIX denoising^77^, which denoises residual subject motion, physiological motions, and artifacts, to the HCP minimally preprocessed task fMRI data. ICA-FIX was implemented via the Quantitative Neuroimaging Environment & Toolbox (QuNex)^78^. For movie-watching fMRI data, we used the FIX-denoised data already provided by the HCP and removed non-steady frames (i.e., the first 15 frames).

### Tensor Independent Component Analysis (tICA)

tICA is a well-validated^25,46^ unsupervised learning technique that decomposes the task fMRI data into a set of brain modes, temporal courses, and subject loadings that reflect the full range of brain activity during a task, from spontaneous activity during resting conditions to task onsets to task maintenance. It can therefore identify, trace the moment-by-moment activity of, and quantify individual differences in task-evoked brain network activity. We applied tICA to the denoised fMRI data from each task separately (and separately for Group 1 and Group 2) using MELODIC^46,79^ in FSL (version 6.0). This identified task-recruited brain networks, including their voxel-wise spatial maps, their frame-wise time courses, and subject loadings (i.e., the extent to which a subject’s data loads on the spatial and temporal pattern of the group-level network) on each network, for each task in both groups. For each task, we ran tICA with different number of components or brain modes estimated (i.e., at varying model orders) to identify the optimal model order for analysis. Because the goal was to investigate large-scale canonical brain networks, the model order of the group ICA needed to be relatively low to result in a coarse spatial scale parcellation of the brain into networks^80^. Model orders that are too high produce finer spatial scale parcellations that reflect excessive segregation of known large-scale networks into individual or bilateral brain areas rather than networks. At the same time, model orders that are too low fail to fully disaggregate known canonical brain networks. Therefore, we tested a range of model orders from 5 to 25 to identify which decomposition parcellated the brain into canonical network patterns while effectively unmixing any noise/artifacts into separate components. For each model order, we visually inspected each component, including the time course and its general linear model fit for the task paradigm, power spectrum and the spatial map to identify components most likely to reflect task-related brain activity (e.g., components with spatial patterns resembling large-scale brain networks assessed both visually and by statistical testing using the Network Correspondence Toolbox^50^, and with temporal courses reflecting one or more aspects of the task time course). Ultimately, we identified to d = 20 to be the optimal model order in our data that best achieved the balance between disaggregating large-scale canonical networks and not producing excessive segregation; this model order is also in line with previous reports examining the correspondence between task activation networks and canonical resting state brain networks^81,82^. Among the 20 components, those identified as noise (e.g., physiological, motion, scanner artifacts, drift) via examination of their time course, power spectrum and spatial distribution were removed from analysis, in line with best practices for identification of ICA noise components^83^. The identified network components were assessed for both spatial and temporal reproducibility across Group 1 and Group 2. Finally, for each group, before input into hierarchical clustering for subject subtyping, subject loadings were normalized by subtracting network loading means and then dividing by network loading standard deviations, on a per network basis.

### Assessing Overlap with Canonical Networks

To further characterize the spatial maps of the tICA-identified network families, we used the network correspondence toolbox (NCT)^50^ to compute overlap (Dice coefficients) between the positive and negative components of each task network and each of 17 canonical large-scale networks defined in the Gordon et al. parcellation^49^. Positive and negative components were analyzed separately because we observed visually that the positive and negative parts of each component typically represented distinct networks. More generally, the tensor ICA identifies brain states or modes comprised of anti-correlated network patterns that have been reported by others during both resting state and task performance (see for example Li et al. 2021^84^) as an important characteristic of brain organization and network dynamics. As such, the positive and negative parts were assessed separately to determine their distinct network makeups. In order to determine statistical significance, a spin test implemented in Python was performed with 1000 permutations for each reference atlas network^50^. Significant overlap was thresholded at *p* < .05.

### Hierarchical Factor Analysis (hFA)

Prior work using hFA identified sets of highly correlated variables from the 87 measured by HCP-YA assessments^29^. These variables included behavioral measures of motor and sensory functioning, alertness, and cognition, including in-scanner task performance measures, as well as self-report measures of emotion, personality, substance use, psychiatric symptoms and general life function^29^. Full details of the procedure are outlined in Schottner et al., 2023^29^. We implemented the Schottner hFA pipeline (https://github.com/connectomicslab/hcp-behavioral-domains) to generate subject factors scores using their five-factor solution which included domains for well-being/positive affect, internalizing/negative affect, processing speed, cognition and substance use^29^. Briefly, prior to the application of factor analysis, missing assessment values were first imputed using Multiple Imputation by Chained Equations. Next, age and gender were regressed out by entering them as predictors in a linear regression. The residuals of this regression served as the inputs to the factor analysis. Reaction times and other error measures were inverted, and data were z-scored across subjects.

### Machine Learning Prediction of Mental Health Scores

For the machine learning regression analyses to predict mental health from across-task and across-network tICA loadings (**Fig. 4**), the number of trees in the random forest was set to 1000. No maximum depth was specified for trees, the minimum number of samples required at each leaf was set to 1, the minimum number of samples required to split an internal node was set to 2, and the maximum number of features to consider when looking for the best split was set to 1.

This analysis was carried out using scikit-learn^85^.

### Hierarchical Clustering

Hierarchical clustering was used for two applications. First, to cluster spatially similar families of brain networks identified from the application of tICA to the five fMRI paradigms (**Fig. 1**). Second, to identify subject subtypes based on brain function and mental health data (**Fig. S1**; **Figs. 5-6**). Hierarchical clustering was implemented by the nets_hierarchy function in the FSLNets package (fsl.fmrib.ox.ac.uk/fsl/fslwiki/FSLNets) using default settings.

## Data Availability

The HCP-YA dataset is publicly available on an open access repository (https://balsa.wustl.edu/), which can be accessed after signing a data use agreement. Individual network maps identified by tICA for the Emotion, Gambling, Working Memory, Social, and Movie-Watching paradigms have been made viewable and accessible in a “Task-Evoked Network Atlas” at https://identifiers.org/neurovault.collection:20820.

## Code Availability

All code used in this paper is freely and publicly available. Code to run tensor independent component analysis (tICA) is available in Functional Magnetic Resonance Imaging of the Brain (FMRIB) Software Library (FSL). Code to run the Network Correspondence Toolbox (NCT) is available here: https://github.com/rubykong/cbig_network_correspondence. Code to run factor analysis on the HCP-YA dataset to generate latent mental health factor scores is available here: https://github.com/connectomicslab/hcp-behavioral-domains. Code to run SHapely Additive exPlanations (SHAP) is available here: https://shap.readthedocs.io/en/latest/. Code to run random forest machine learning regression is available here: https://scikit-learn.org/stable/modules/generated/sklearn.ensemble.RandomForestClassifier.html.

## Declaration of Interests

The authors declare no competing interests.

## Acknowledgements

Work funded by the National Institutes of Aging 1RF1AG078304-01 to LN and DH.

---

**Supplementary Table S1.**
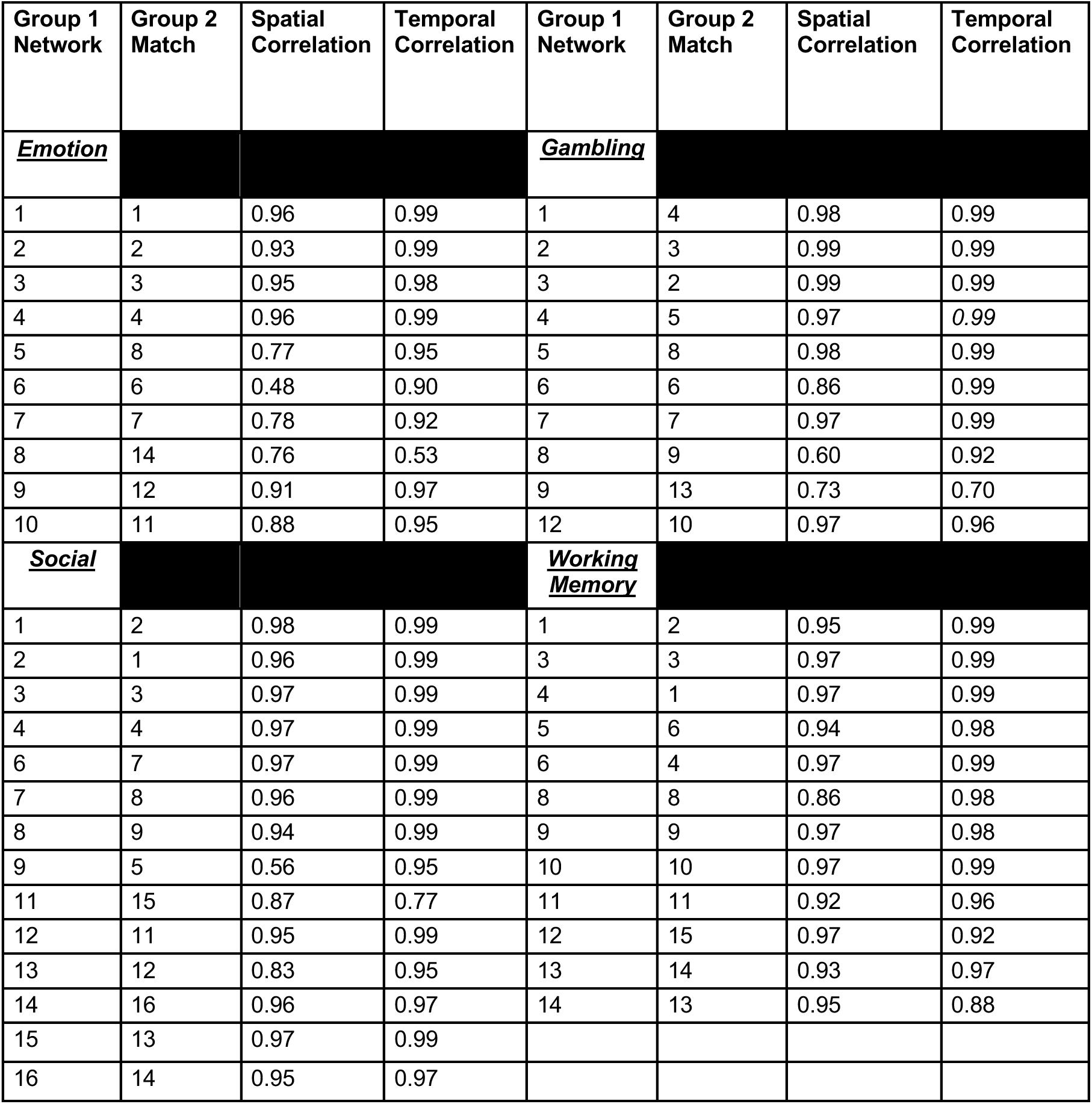
Spatiotemporal correspondence of 46 task-evoked networks that replicate across Group 1 (discovery sample) and Group 2 (replication sample).

**Supplementary Figure S1.**
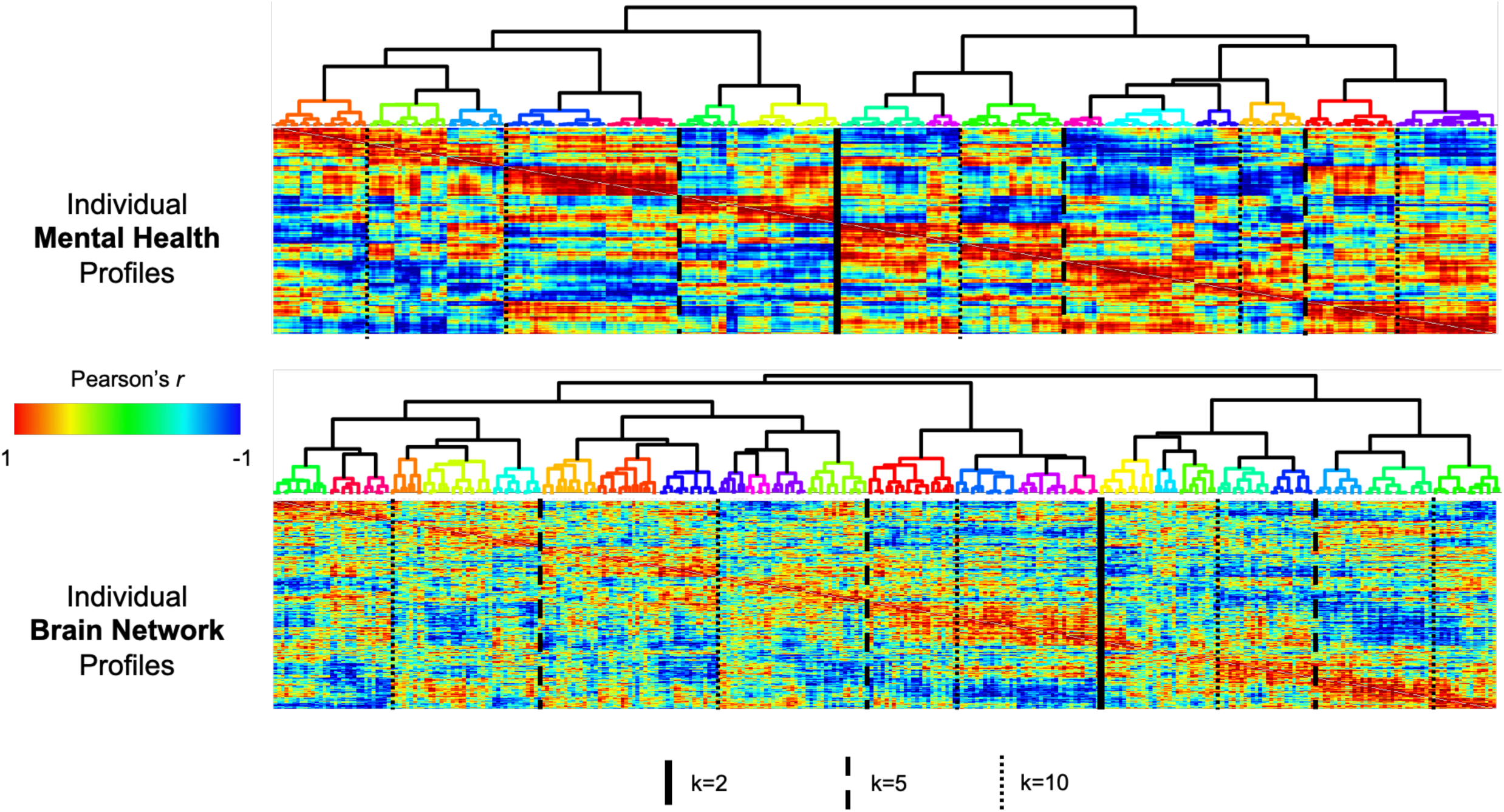
Hierarchical clustering of subjects into subtypes based on their mental health profiles (top) and based on their brain function profiles (bottom). Correlation matrices quantify the similarity between each pair of subjects. Starting from the top of each clustering tree, we quantified the number of branches at each successive vertical level of the tree. When a vertical level was present in one portion of the tree but not the others, the total number of branches at that level was counted as the number of new branches from the portion with the new level plus the number of branches from the previous level in the remainder of the tree. The mental health and brain function trees shared an equivalent number of branches at three levels, where the number of groups k was k=2, k=5 and k=10. At the finest clustering level, 16 mental health subtypes and 25 brain function subtypes were identified.

**Supplementary Figure S2.**
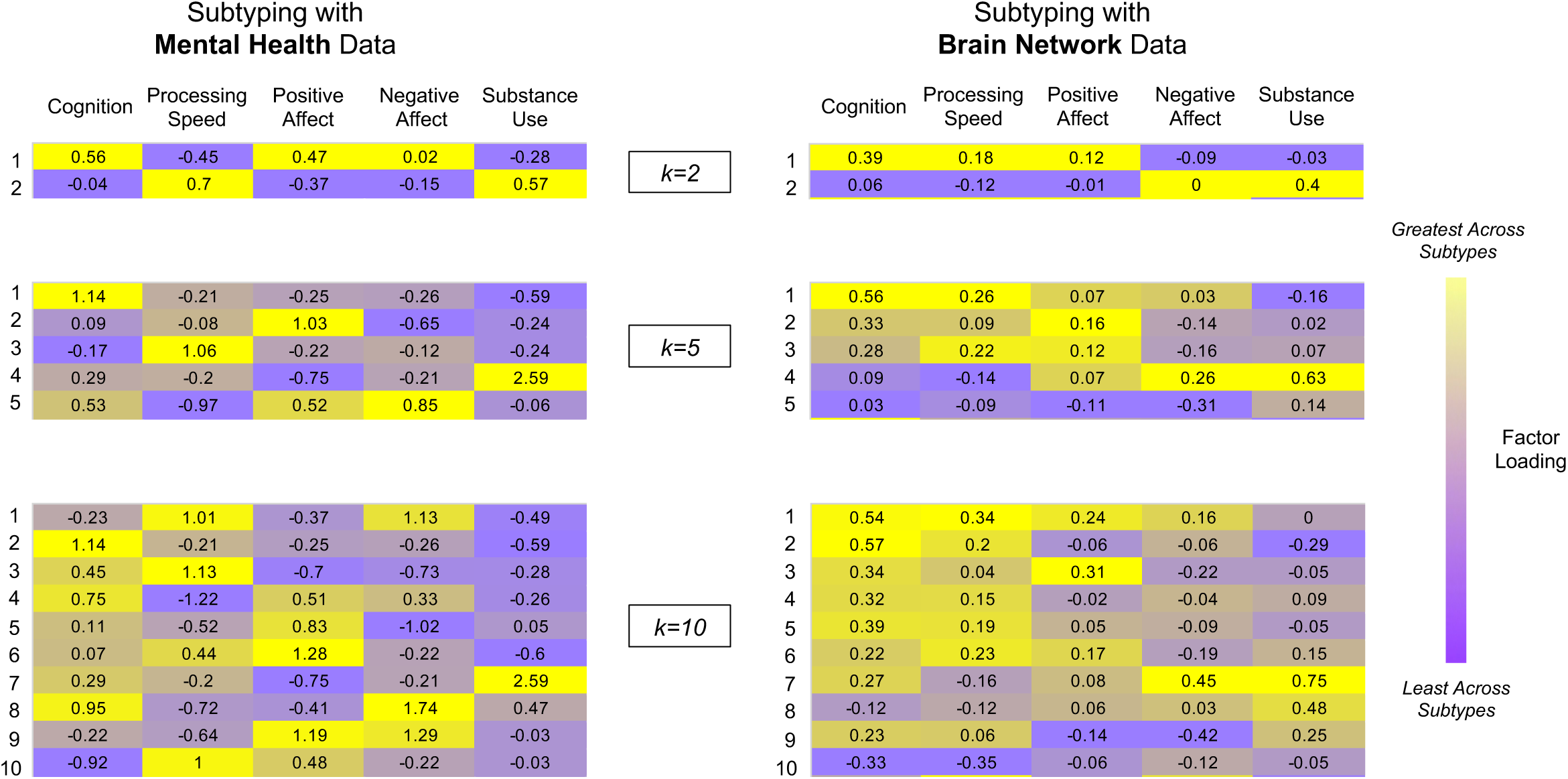
Average mental health profile of each mental health-based subtype and each brain function-based subtype. Side-by-side comparisons of the mean mental health factor scores of each mental health subtype and its most closely corresponding brain function subtype. Correspondence was operationalized by determining the mental health–brain function subtype pairs with the maximally correlated mean mental health factor score profiles. Within each clustering level (i.e., k=2, 5 or 10) color-coding is applied on a column-wise basis to highlight the relative scoring of each subtype on each mental health factor. Subtyping subjects via their mental health profile revealed the following groups. At the broadest (k=2) level, the sample split into a group with higher cognition and positive affect, and lower processing speed and substance use (subtype 1); and a group with lower cognition and positive affect, and higher processing speed and substance use (subtype 2). Negative affect was roughly similar across the two groups. At the k=5 level, subjects with the highest average cognition had the lowest average substance use (subtype 1); subjects with the highest average positive affect had the lowest average negative affect (subtype 2); subjects with the highest average processing speed had the lowest average cognition scores (subtype 3); subjects with the highest average substance use had the lowest average positive affect (subtype 4); and subjects with the highest average negative affect had the lowest average processing speed (subtype 5). At the k=10 level, subjects with high substance use levels were distinguished by those with very low positive affect (subtype 7) and those with very high negative affect and very high cognition (subtype 8). Similarly, three types of very high cognition individual were identified: those with low negative affect and very low substance use (subtype 2), those with high positive affect and low substance use (subtype 4), and those with very high negative affect and high substance use. Interestingly, one subtype was defined by both very high positive and negative affect (subtype 9).

**Supplementary Figure S3.**
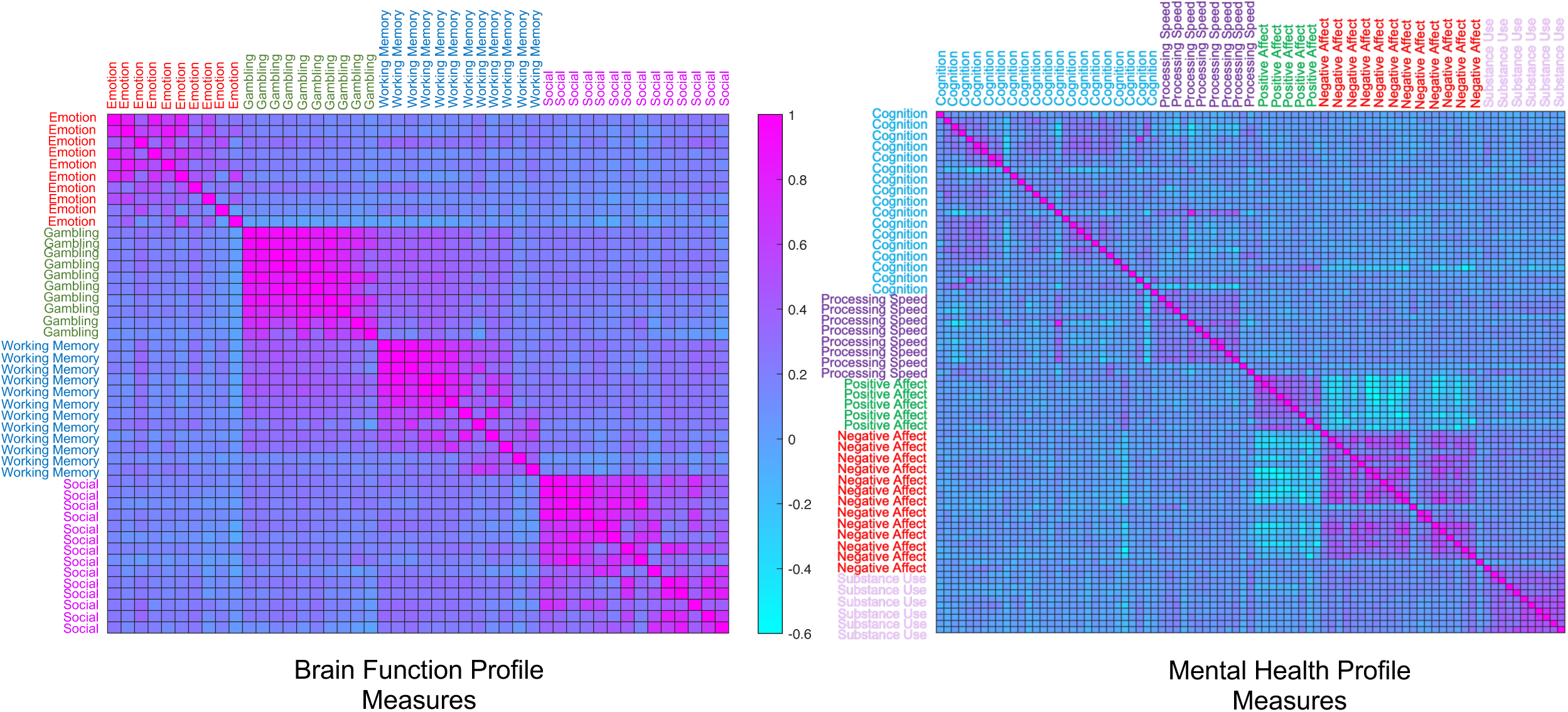
Diversity of brain function and mental health assessments. Magnitudes by which subject scores on each measurement item correlated with subject scores on every other measurement item, separately, for brain function and mental health data. For brain function, measurement items included activation strengths on the 10 Emotion task-evoked networks, 10 Gambling task-evoked networks, 12 Working Memory task-evoked networks, and 14 Social task-evoked networks that replicated across groups. For mental health profiles, measurement items included factor loadings on composites of 25 measures of cognition, 21 measures of negative affect/internalizing, 10 measures of positive affect/well-being, 18 measures of processing speed, and 12 measures of substance use. Correlation matrices were computed on data from the 324 subjects from Group 1 with all fMRI task data.

## Notes

https://identifiers.org/neurovault.collection:20820

### Competing Interest Statement

The authors have declared no competing interest.

### Summary of Updates

Additional analyses have been added (Figures 5-7)

