## Supplemental Material for "Weak brain-mental health associations in population neuroimaging tasks reflect a signal evocation bottleneck"

| Group 1 Network | Group 2 Match | Spatial Correlation | Temporal Correlation | Group 1 Network | Group 2 Match | Spatial Correlation | Temporal Correlation |
|---|---|---|---|---|---|---|---|
| ***Emotion*** | | | | ***Gambling*** | | | |
| 1 | 1 | 0.96 | 0.99 | 1 | 4 | 0.98 | 0.99 |
| 2 | 2 | 0.93 | 0.99 | 2 | 3 | 0.99 | 0.99 |
| 3 | 3 | 0.95 | 0.98 | 3 | 2 | 0.99 | 0.99 |
| 4 | 4 | 0.96 | 0.99 | 4 | 5 | 0.97 | *0.99* |
| 5 | 8 | 0.77 | 0.95 | 5 | 8 | 0.98 | 0.99 |
| 6 | 6 | 0.48 | 0.90 | 6 | 6 | 0.86 | 0.99 |
| 7 | 7 | 0.78 | 0.92 | 7 | 7 | 0.97 | 0.99 |
| 8 | 14 | 0.76 | 0.53 | 8 | 9 | 0.60 | 0.92 |
| 9 | 12 | 0.91 | 0.97 | 9 | 13 | 0.73 | 0.70 |
| 10 | 11 | 0.88 | 0.95 | 12 | 10 | 0.97 | 0.96 |
| ***Social*** | | | | ***Working Memory*** | | | |
| 1 | 2 | 0.98 | 0.99 | 1 | 2 | 0.95 | 0.99 |
| 2 | 1 | 0.96 | 0.99 | 3 | 3 | 0.97 | 0.99 |
| 3 | 3 | 0.97 | 0.99 | 4 | 1 | 0.97 | 0.99 |
| 4 | 4 | 0.97 | 0.99 | 5 | 6 | 0.94 | 0.98 |
| 6 | 7 | 0.97 | 0.99 | 6 | 4 | 0.97 | 0.99 |
| 7 | 8 | 0.96 | 0.99 | 8 | 8 | 0.86 | 0.98 |
| 8 | 9 | 0.94 | 0.99 | 9 | 9 | 0.97 | 0.98 |
| 9 | 5 | 0.56 | 0.95 | 10 | 10 | 0.97 | 0.99 |
| 11 | 15 | 0.87 | 0.77 | 11 | 11 | 0.92 | 0.96 |
| 12 | 11 | 0.95 | 0.99 | 12 | 15 | 0.97 | 0.92 |
| 13 | 12 | 0.83 | 0.95 | 13 | 14 | 0.93 | 0.97 |
| 14 | 16 | 0.96 | 0.97 | 14 | 13 | 0.95 | 0.88 |
| 15 | 13 | 0.97 | 0.99 | | | | |
| 16 | 14 | 0.95 | 0.97 | | | | |

**Supplementary Table S1.** Spatiotemporal correspondence of 46 task-evoked networks that replicate across Group 1 (discovery sample) and Group 2 (replication sample).

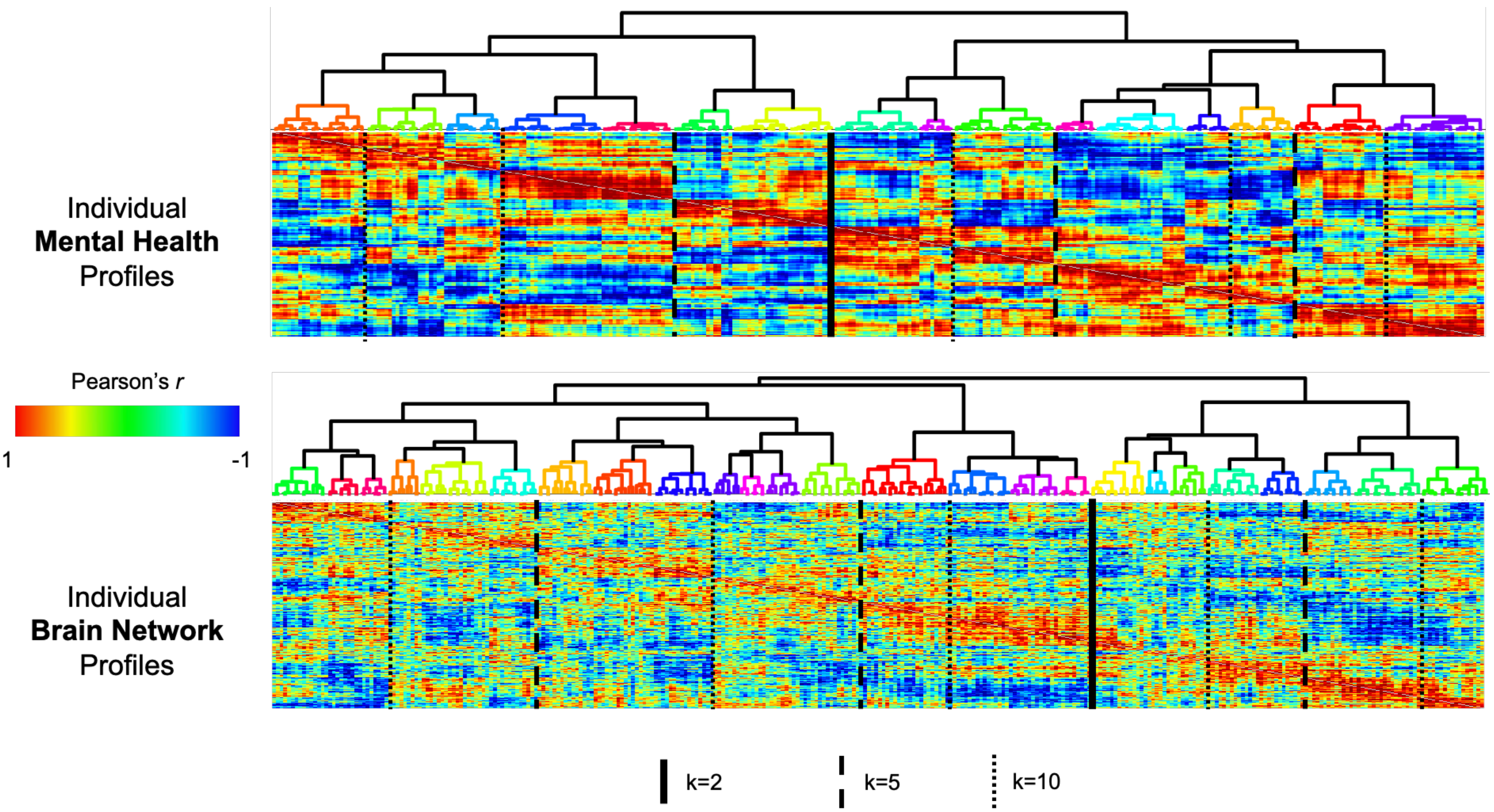


**Supplementary Figure S1.** *Hierarchical clustering of subjects into subtypes based on their mental health profiles (top) and based on their brain function profiles (bottom).* Correlation matrices quantify the similarity between each pair of subjects. Starting from the top of each clustering tree, we quantified the number of branches at each successive vertical level of the tree. When a vertical level was present in one portion of the tree but not the others, the total number of branches at that level was counted as the number of new branches from the portion with the new level plus the number of branches from the previous level in the remainder of the tree. The mental health and brain function trees shared an equivalent number of branches at three levels, where the number of groups k was k=2, k=5 and k=10. At the finest clustering level, 16 mental health subtypes and 25 brain function subtypes were identified.

Subtyping with **Mental Health** Data

Subtyping with **Brain Network** Data

*k=2*

Mental Health Data:

| | Cognition | Processing Speed | Positive Affect | Negative Affect | Substance Use |
|---|---|---|---|---|---|
| 1 | 0.56 | -0.45 | 0.47 | 0.02 | -0.28 |
| 2 | -0.04 | 0.7 | -0.37 | -0.15 | 0.57 |

Brain Network Data:

| | Cognition | Processing Speed | Positive Affect | Negative Affect | Substance Use |
|---|---|---|---|---|---|
| 1 | 0.39 | 0.18 | 0.12 | -0.09 | -0.03 |
| 2 | 0.06 | -0.12 | -0.01 | 0 | 0.4 |

*k=5*

Mental Health Data:

| | Cognition | Processing Speed | Positive Affect | Negative Affect | Substance Use |
|---|---|---|---|---|---|
| 1 | 1.14 | -0.21 | -0.25 | -0.26 | -0.59 |
| 2 | 0.09 | -0.08 | 1.03 | -0.65 | -0.24 |
| 3 | -0.17 | 1.06 | -0.22 | -0.12 | -0.24 |
| 4 | 0.29 | -0.2 | -0.75 | -0.21 | 2.59 |
| 5 | 0.53 | -0.97 | 0.52 | 0.85 | -0.06 |

Brain Network Data:

| | Cognition | Processing Speed | Positive Affect | Negative Affect | Substance Use |
|---|---|---|---|---|---|
| 1 | 0.56 | 0.26 | 0.07 | 0.03 | -0.16 |
| 2 | 0.33 | 0.09 | 0.16 | -0.14 | 0.02 |
| 3 | 0.28 | 0.22 | 0.12 | -0.16 | 0.07 |
| 4 | 0.09 | -0.14 | 0.07 | 0.26 | 0.63 |
| 5 | 0.03 | -0.09 | -0.11 | -0.31 | 0.14 |

*k=10*

Mental Health Data:

| | Cognition | Processing Speed | Positive Affect | Negative Affect | Substance Use |
|---|---|---|---|---|---|
| 1 | -0.23 | 1.01 | -0.37 | 1.13 | -0.49 |
| 2 | 1.14 | -0.21 | -0.25 | -0.26 | -0.59 |
| 3 | 0.45 | 1.13 | -0.7 | -0.73 | -0.28 |
| 4 | 0.75 | -1.22 | 0.51 | 0.33 | -0.26 |
| 5 | 0.11 | -0.52 | 0.83 | -1.02 | 0.05 |
| 6 | 0.07 | 0.44 | 1.28 | -0.22 | -0.6 |
| 7 | 0.29 | -0.2 | -0.75 | -0.21 | 2.59 |
| 8 | 0.95 | -0.72 | -0.41 | 1.74 | 0.47 |
| 9 | -0.22 | -0.64 | 1.19 | 1.29 | -0.03 |
| 10 | -0.92 | 1 | 0.48 | -0.22 | -0.03 |

Brain Network Data:

| | Cognition | Processing Speed | Positive Affect | Negative Affect | Substance Use |
|---|---|---|---|---|---|
| 1 | 0.54 | 0.34 | 0.24 | 0.16 | 0 |
| 2 | 0.57 | 0.2 | -0.06 | -0.06 | -0.29 |
| 3 | 0.34 | 0.04 | 0.31 | -0.22 | -0.05 |
| 4 | 0.32 | 0.15 | -0.02 | -0.04 | 0.09 |
| 5 | 0.39 | 0.19 | 0.05 | -0.09 | -0.05 |
| 6 | 0.22 | 0.23 | 0.17 | -0.19 | 0.15 |
| 7 | 0.27 | -0.16 | 0.08 | 0.45 | 0.75 |
| 8 | -0.12 | -0.12 | 0.06 | 0.03 | 0.48 |
| 9 | 0.23 | 0.06 | -0.14 | -0.42 | 0.25 |
| 10 | -0.33 | -0.35 | -0.06 | -0.12 | -0.05 |

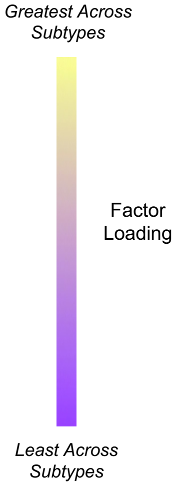


**Supplementary Figure S2.** *Average mental health profile of each mental health-based subtype and each brain function-based subtype.* Side-by-side comparisons of the mean mental health factor scores of each mental health subtype and its most closely corresponding brain function subtype. Correspondence was operationalized by determining the mental health–brain function subtype pairs with the maximally correlated mean mental health factor score profiles. Within each clustering level (i.e., k=2, 5 or 10) color-coding is applied on a column-wise basis to highlight the relative scoring of each subtype on each mental health factor. Subtyping subjects via their mental health profile revealed the following groups. At the broadest (k=2) level, the sample split into a group with higher cognition and positive affect, and lower processing speed and substance use (subtype 1); and a group with lower cognition and positive affect, and higher processing speed and substance use (subtype 2). Negative affect was roughly similar across the two groups. At the k=5 level, subjects with the highest average cognition had the lowest average substance use (subtype 1); subjects with the highest average positive affect had the lowest average negative affect (subtype 2); subjects with the highest average processing speed had the lowest average cognition scores (subtype 3); subjects with the highest average substance use had the lowest average positive affect (subtype 4); and subjects with the highest average negative affect had the lowest average processing speed (subtype 5). At the k=10 level, subjects with high substance use levels were distinguished by those with very low positive affect (subtype 7) and those with very high negative affect and very high cognition (subtype 8). Similarly, three types of very high cognition individual were identified: those with low negative affect and very low substance use (subtype 2), those with high positive affect and low substance use (subtype 4), and those with very high negative affect and high substance use. Interestingly, one subtype was defined by both very high positive and negative affect (subtype 9).

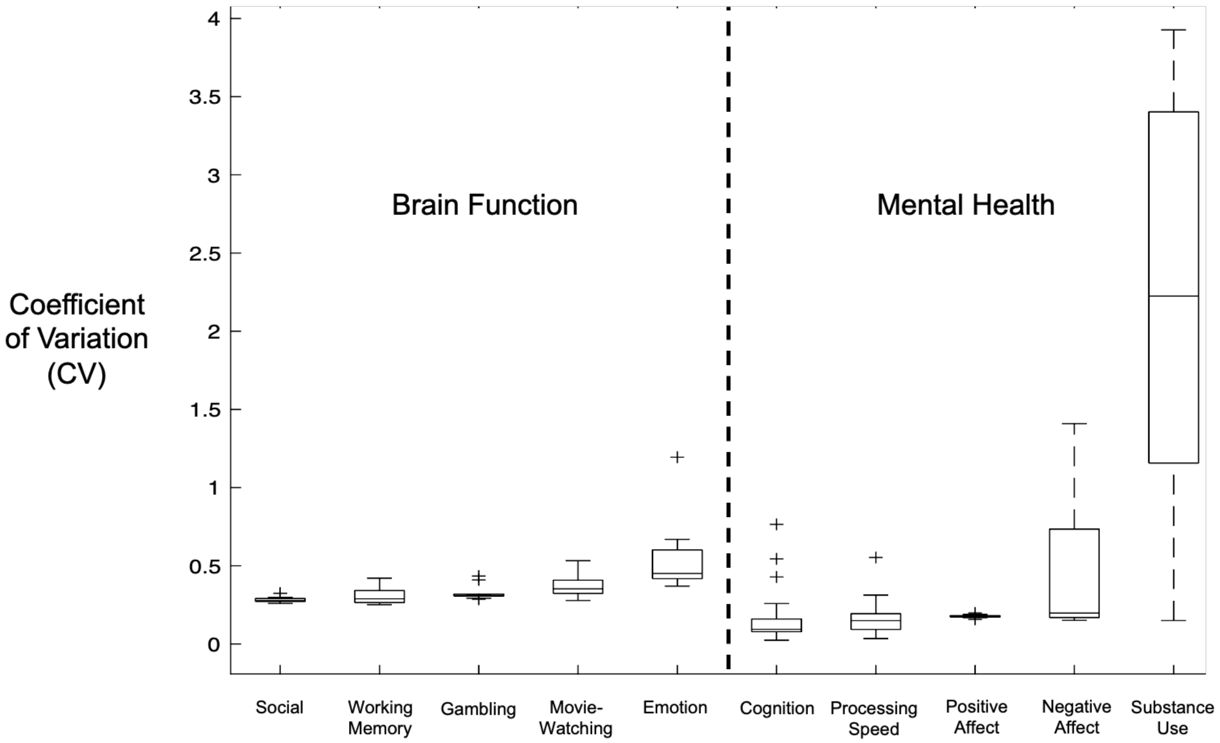


**Supplementary Figure S3.** *Interindividual variability of brain function and mental health metrics.* Coefficients of variation (i.e., standard deviation/mean) were computed for each of the 77 task-evoked brain networks and each of the 87 mental health assessments. Plotted are the distributions (minimum, first quartile, median, third quartile, maximum, outliers) of CVs for metrics in each category.

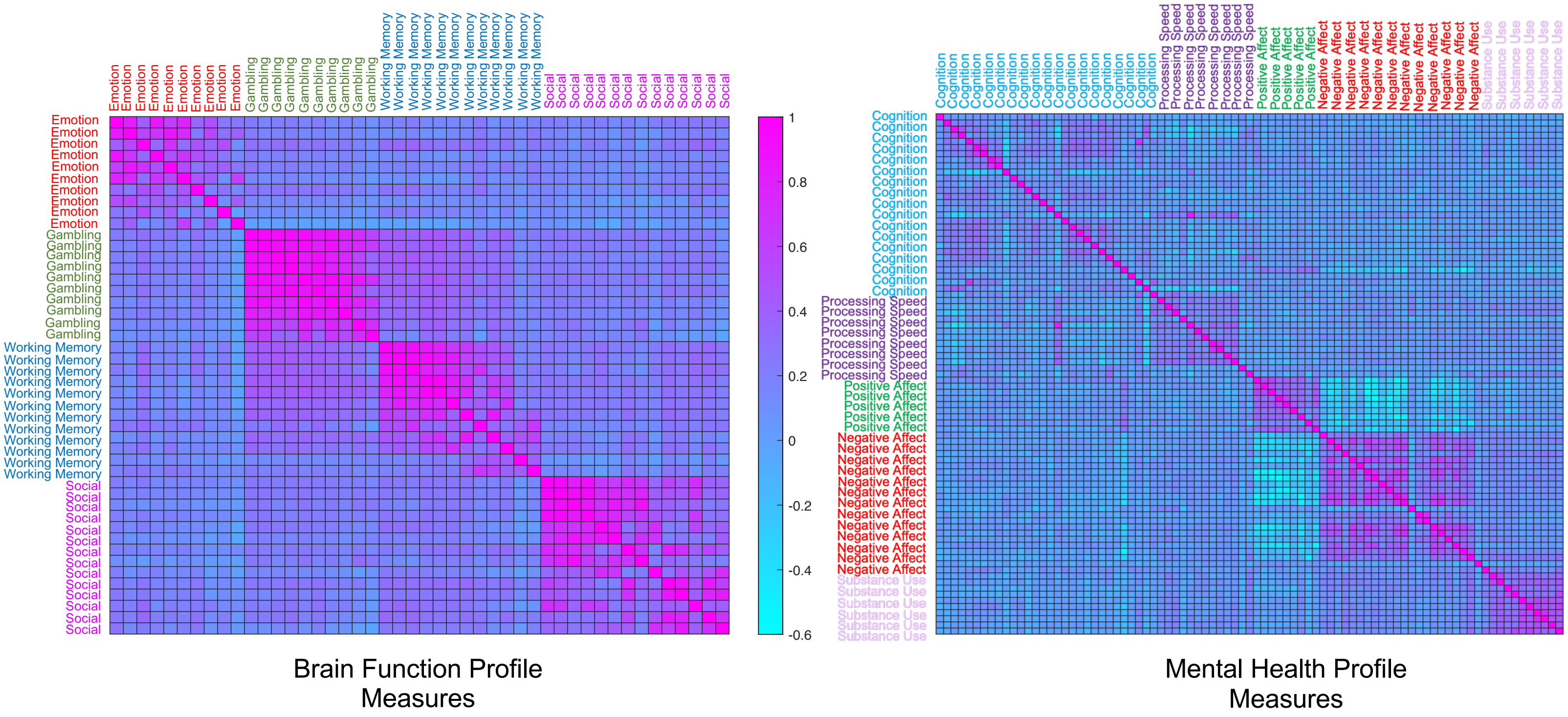


**Supplementary Figure S4.** *Diversity of brain function and mental health assessments.* Magnitudes by which subject scores on each measurement item correlated with subject scores on every other measurement item, separately, for brain function and mental health data. For brain function, measurement items included activation strengths on the 10 Emotion task-evoked networks, 10 Gambling task-evoked networks, 12 Working Memory task-evoked networks, and 14 Social task-evoked networks that replicated across groups. For mental health profiles, measurement items included factor loadings on composites of 25 measures of cognition, 21 measures of negative affect/internalizing, 10 measures of positive affect/well-being, 18 measures of processing speed, and 12 measures of substance use. Correlation matrices were computed on data from the 324 subjects from Group 1 with all fMRI task data.
